# Evolution of chemosensory tissues and cells across ecologically diverse *Drosophilids*

**DOI:** 10.1101/2023.04.14.536691

**Authors:** Gwénaëlle Bontonou, Bastien Saint-Leandre, Tane Kafle, Tess Baticle, Afrah Hassan, Juan Antonio Sánchez-Alcañiz, Roman J. Arguello

## Abstract

Chemosensory systems display exceptional variation between species, but little is known about how the evolution of gene expression and cell types contribute to this diversity. We have generated transcriptomes for five chemosensory tissues across six ecologically diverse *Drosophila* species and integrated their analyses with single-cell datasets to address these questions. The evolution of chemosensory transcriptomes has been predominantly shaped by stabilizing selection, but several thousand genes have nevertheless evolved expression changes in each tissue. Phylogenetic analyses of differentially expressed genes revealed strong evidence that their expression changes have been driven by a combination of *cis*-regulatory and cell composition evolution. We have also found that chemosensory-related gene families have undergone pervasive expression level changes and numerous species-specific expression gains/losses. Follow-up experiments revealed several chemoreceptors that evolved novel patterns of tissue and cellular expression that likely contribute to sensory differences. Finally, analyses of the genes that are differentially expressed between sexes uncovered extensive species-specific differences. Among these rapid changes, we discovered a *D. melanogaster-*specific excess of male-biased gene expression in its forelegs and identified sensory and muscle cells as the primary source of this dimorphism. Together, our analyses provide new insights for understanding evolutionary changes in ecologically key tissues at both global and individual gene levels.

## Introduction

The abilities of animals to perceive their chemical environments are remarkably variable. Chemosensory receptor protein families and the cell types where they are expressed have multiple evolutionary origins^1–6^, and the tissues that contain them can differ dramatically across species in morphology and anatomical distributions^7–12^. For example, while taste perception in mammals is largely restricted to gustatory cells located in the mouth - and primarily the tongue, aquatic vertebrates have taste cells distributed externally on their skin^13–17^. Insects have evolutionarily distinct taste receptors and cells that are also broadly distributed across their bodies, including their mouth parts, legs, wing margins, and ovipositors^18^. Appendages involved in smell are generally more restricted to animals’ heads but also differ dramatically among taxa, as exemplified by the bulbous nose of the Proboscis Monkey or the feathery antennae of moths. In addition to differences among species, striking evolutionary changes have also arisen between sexes within species. Sexual dimorphisms in chemosensory perception and organ morphology often evolve rapidly and have been attributed to differences in sex-specific physiological states, sexual selection, and sex-specific nutritional needs, among other factors^19–22^.

Understanding the molecular basis of chemosensory evolution is important for fundamental and applied biology. Insights into the genes and expression changes that underlie species’ chemosensory differences help us understand how nervous systems adapt in response to varying ecologies and provide the basis for managing disease vectors and agricultural pests. For instance, research on insect chemosensation has advanced our understanding of how mosquitoes track human odors, with important implications for human health^23–25^, and has aided in the development of novel farming methods that reduce crop infestation^26^. While these applications draw on knowledge of chemosensation from a broad range of biological models, much of what we know derives from research on *Drosophila melanogaster*.

Research on *D. melanogaster* has led to extensive knowledge about the development of its nervous system and chemosensory appendages and has generated a nearly complete mapping of its full set of olfactory and gustatory receptor proteins to specific neuron populations. This work has provided the basis for many pioneering functional and behavioral studies^27–32^. In addition, advances in connectomics and single-cell transcriptomics applied to *D. melanogaster*’s nervous system are helping to identify new developmental factors, describe cellular diversity in chemosensory tissues, and characterize synapse-level connections from the peripheral chemosensory neurons to the central brain^33–41^. Beyond its role as a preeminent model for chemosensory biology, *D. melanogaster* and its closely-related species have also long been a model system for evolutionary genetics and speciation^42–47^. The phylogenetic relationships among the *D. melanogaster* species group are well-resolved and include lineages of diverse ages and ecologies. This system, therefore, provides a valuable opportunity to ask how evolutionary forces and environments shape chemosensory systems^22, 48, 49^. However, beyond the meticulous molecular and cellular characterization of *D. melanogaster*’s chemosensory tissues, little is known about how they evolve between species.

To address this question, we have carried out a comparative transcriptomic experiment in which we generated bulk RNA-sequencing (RNA-seq) datasets for five chemosensory tissues: larva head (mixed sex), ovipositor (female), forelegs (female and male), antenna (female and male), and proboscis with maxillary palps (female and male). These samples were collected from six ecologically diverse species in the *D. melanogaster* species group that share common ancestors between ∼0.25-15 million years ago^44, 50–53^: *D. melanogaster*, *D. sechellia*, *D. simulans*, *D. santomea*, *D. erecta*, and *D. suzukii* (Fig. 1A). *D. sechellia*, is endemic to Seychelles and an extreme specialist on the fruit of *Morinda citrifolia,* which is toxic to the other species^54^. *D. santomea* is endemic to São Tomé and adapted to high elevation mist forests^55, 56^. *D. erecta* is restricted to west-central Africa and is thought to be an opportunistic specialist on the fruits of *Pandanus*^57^. *D. suzukii* originated in Eastern Asia but has expanded rapidly worldwide in the last decade^58, 59^. Unlike the other species, *D. suzukii* females oviposit in ripe soft-bodied fruits and, as a result, have become a global agricultural pest^60–65^. Both *D. simulans* and *D. melanogaster* are generalists that feed on a broad range of decaying fruits and have nearly worldwide distributions^66^. Using these data, we have asked how different evolutionary histories and ecological specializations have driven rates of transcriptomic change across chemosensory tissues and how pleiotropy has shaped gene expression profiles. Given the importance of chemosensory perception in *Drosophila*’s mating systems and sexually dimorphic behaviors, we have also asked how expression differences between the sexes have evolved. These data can be explored with our dashboard available at: https://evoneuro.shinyapps.io/ctct/.

**Fig. 1.**
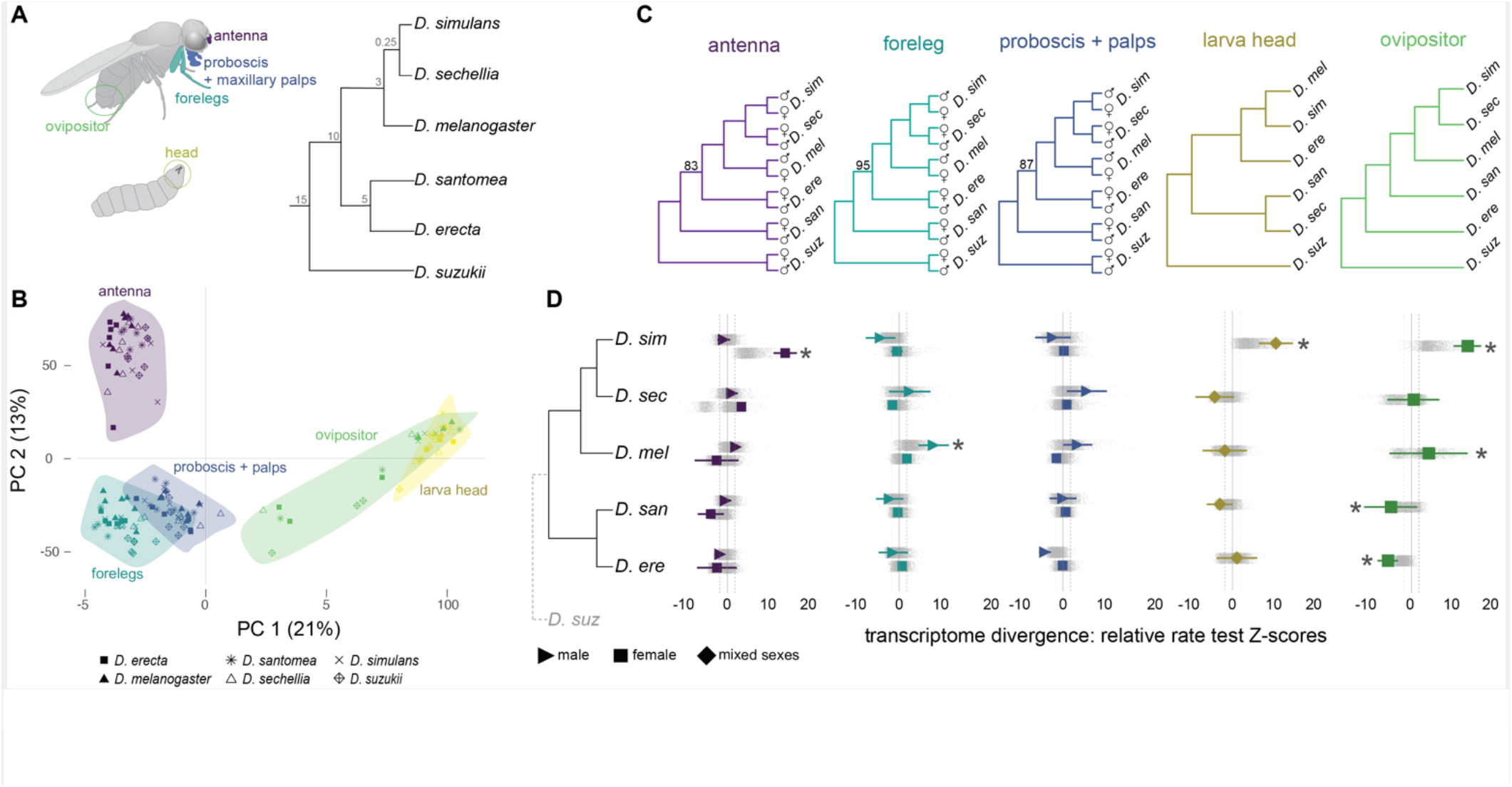
Chemosensory tissue transcriptome evolution. **(A)** Overview of the chemosensory tissues (left) and species (right) used in this study. The numbers at the nodes of the species tree are the estimated divergence dates in millions of years. **(B)** PCA of the transcriptomic datasets using 1:1 orthologs. The percentages on the axes are the amount of variation explained by the PCs. **(C)** Clustering of the transcriptomic datasets (1:1 orthologs) according to species and sex. Species names are abbreviated to the first three letters. **(D)** Relative rate test results arranged by the species phylogeny. Colored shapes and lines display the mean and standard deviation of Z-scores from the full set of 1:1 orthologs. *D. suzukii*, noted with the dashed line and gray font, was used as the outgroup species. Gray data points are Z-scores that resulted from repeating the tests with subsampled datasets (Methods). Asterisks denote the significantly elevated (positive values) or reduced (negative values) rates of gene expression change (Wilcoxon test comparing Z-score distribution to the minimum and maximum values of non-significant Z-scores: dotted lines). Species names are abbreviated to the first three letters.

## Results

### Relationships between sensory tissue transcriptomes

To study the evolution of gene expression in the main chemosensory tissues of closely-related *Drosophila*, we generated bulk RNA-seq datasets for six ecologically diverse species and five sensory tissues (Fig. 1A; Methods). On average, we obtained 43 million mapped reads per sample with high correlations across triplicates (average Pearson correlation coefficient = 0.98). To overcome annotation biases, we used these datasets to produce equivalent *de novo* gene annotations and used the resulting gene sets for orthology/paralogy assignments. This approach resulted in similar genome annotations with BUSCO scores ranging from 91.9 - 97.3%, indicating a well-balanced dataset for cross-species comparisons.

We began investigating the relationship between chemosensory tissue transcriptomes by conducting a principal component (PC) analysis on expression levels of ∼12,000 genes with a one-to-one relationship across the six species in our sample (1:1 orthologs; Fig. 1B). The first principle component (PC1) separates the three appendage samples from the larval head and ovipositor samples. The genes that contribute the most to the negative loading of PC1 are enriched for gene ontology (GO) terms related to cilia, cell projections/axons, and synapses, among other neural categories. These terms contrast with the enrichment of cell cycle, organelle, and nucleus-related terms that most contribute to the top positive loadings of PC1 (Fig. S1). Analysis of correlated expression changes across multiple genes also identified appendage-specific expression modules that are enriched for cilium, dendrite, and chemosensory terms that load negatively on PC1 and larval/ovipositor-specific modules that are enriched for cell cycle ontology terms that load positively on PC1 (Fig. S1). The second principal component (PC2) separates the antenna from the other samples and is enriched for GO terms related to olfactory, dendrite, and sensory function for the top positive loadings. We again identified an antenna-specific module that is enriched for olfactory receptor, odorant binding, and dendrite terms that load positively on PC2 (Fig. S1). The gene set defining this antenna module negatively correlates with a muscle-related gene module that is enriched in the forelegs and proboscis datasets (Fig. S1), highlighting both neural and structural genes underlying the chemosensory tissue transcriptome differences.

We observed further separation between the antenna, ovipositor, and larva clusters with additional PC pairings, but the foreleg and proboscis+palps transcriptomes always overlap (Fig. S2). This tight pairing indicates that the forelegs and proboscis+palps are more similar to each other than to the antenna. This finding contrasts with the results of developmental genetic analyses of appendage patterning genes that suggested the proboscis was distinct from the legs and antenna - at least at the adult transcriptomic level^68^. Among the five tissues, the ovipositor samples varied the most in PC space, reflecting the lower correlation across some replicates for this tissue (Methods). Despite this variation, the clustering separated the *D. suzukii* samples, for which the replicates were highly correlated (average Pearson correlation coefficient = 0.97). This *D. suzukii* difference is notable because the females of this species differ from the others in their preference for ovipositing in ripening fruits (instead of overripe/rotting fruits) and have evolved an elongated serrated ovipositor that punctures fruit skins^69^.

### Stabilizing selection shapes sensory transcriptomes, but there are exceptions

To investigate the clustering of the transcriptomic datasets on a species level, we estimated expression distances by applying an evolutionary model of transcriptome divergence^70^. This clustering largely recapitulates the known phylogeny (Figs. 1C). For all tissues except the larva head, the consistent difference between the species’ genetic relationships and the transcriptomic clustering is the lack of an internal node shared by *D. erecta* and *D. santomea*. The transcriptomic clustering of the larval head dataset results in additional discrepancies, with *D. erecta* grouping with *D. melanogaster* and *D. simulans*, *D. sechellia* and *D. santomea* grouping together (Fig. 1C). This pattern points to a more complex evolutionary history of gene expression evolution for the larva head compared to the other tissues.

The distinct ecological differences and evolutionary histories among these six species led us to hypothesize that their chemosensory transcriptomes have evolved at different rates. We tested for these differences by applying relative rate tests, which use a pair of ingroup species with an outgroup species to determine whether one of the two ingroup lineages has a significantly elevated rate of transcriptomic change^70^. We applied this test to all 1:1 orthologs for all species-pairs (setting *D. suzukii* as the outgroup) and found that the distribution of test scores (Z-scores) for the majority of the species-tissue comparisons are largely consistent with equal rates of transcriptomic change, indicating that stabilizing selection is the prevailing evolutionary force acting on these tissues (Fig. 1D). We obtained consistent results when examining the distribution of Z-scores based on subsampled sets of the 1:1 orthologs (Fig. 1D).

Despite the predominant role of stabilizing selection, we also identified several tissues that stand out with significantly elevated species-specific and sex-specific rates of transcriptomic evolution. *D. simulans* has a significantly elevated rate of evolution for its female antenna transcriptome (Wilcoxon signed-rank test V = 4.6e+06, *p* < 0.001), larva head transcriptome (Wilcoxon signed-rank test V = 7.1e+06, *p* < 0.001), and ovipositor transcriptome (Wilcoxon signed-rank test V = 2.9e+06, *p* < 0.001). In addition, *D. melanogaster’s* male forelegs and ovipositor transcriptomes have significantly elevated rates of transcriptome change (Wilcoxon signed-rank test V = 6.6e+06, *p* < 0.001 *and* V = 3.1e+06, *p* < 0.001, respectively). In contrast, the ovipositor transcriptomes from *D. santomea* and *D. erecta* were both found to have significantly lower expression divergence compared to the other species (Wilcoxon signed-rank test V = 3.1e+06, *p* < 0.001 and V = 2.3e+06, *p* < 0.001, respectively). Overall, these global analyses of transcriptomic differences highlight a limited set of sensory tissues as rapidly evolving among the species, possibly reflecting key ecological and/or functional differences. They also provide evidence for significant sex differences that exist within species (see below).

### Genes change expression in multiple tissues but at different times

Our finding that stabilizing selection is the principal mode of selection acting on the chemosensory transcriptomes as a whole does not mean that individual genes have not evolved novel expression patterns during the species’ diversification. We thus asked which genes have changed expression among species as well as when in the past the changes have taken place. Using phylogenetically-informed tests for expression changes applied to our set of 1:1 orthologs, we detected several thousand differentially expressed genes for each tissue. Most of these expression changes occurred in only one lineage. The total number of genes for which we detected changes ranges from 8,499 in male antennae to 7,501 in female legs (Fig. 2A, Table S1). Analysis of the functional categories enriched by these differentially expressed genes highlighted combinations of developmental/morphological, neural/sensory, and gene regulation terms, among others, in varying proportions along extant and past lineages (Fig. S3). The elevated number of expression changes identified in *D. simulans* female antenna (1,609) and larva (2,667) and in *D. melanogaster* male forelegs (1,796), confirms a history of punctuated rates of expression evolution for these tissues (Fig. 2A; see also Fig. 1D).

**Fig. 2.**
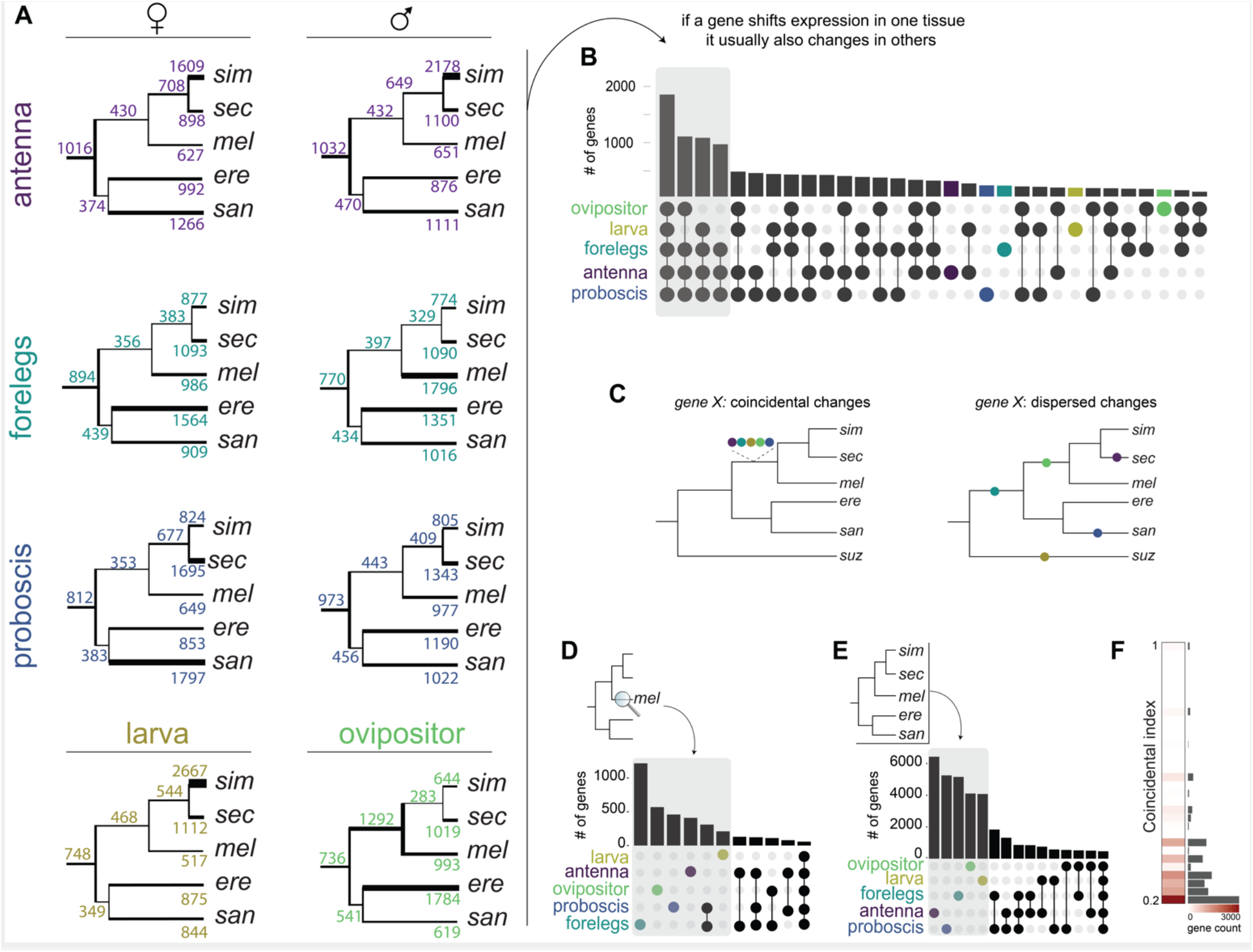
Expression changes over branches and tissues. **(A)** Expression changes inferred across the species’ phylogeny across all tissues. The number above each branch is the total number of expression changes (up and down), and the thickness of the branch is proportional to that number. **(B)** Quantification of the genes that change in expression across multiple tissues. The height of each bar indicates the number of genes that have changed expression across the set of tissues indicated by the darkened/colored circles. **(C)** Schematic illustrating a gene having expression changes involving multiple tissues that were coincidental (occurring on a single branch) or dispersed (occurring across multiple branches). **(D)** Quantification of the coincidental expression changes along the branch leading to *D. melanogaster*. The height of each bar indicates the number of genes that have changed expression across the set of tissues indicated by the darkened/colored circles. The plot was truncated at the bin containing the overlap of the five tissues. **(E)** Quantification of the coincidental expression changes over all branches in the phylogeny. The height of each bar indicates the number of genes that have changed expression across the set of tissues indicated by the darkened/colored circles. The plot was truncated at the bin containing the overlap of the five tissues. **(F)** Summary of the distribution of the “coincidental index” distribution for all expression changes.

Having identified gene expression changes for each tissue on its own, we next questioned how often a given gene changed expression in multiple tissues. Quantifying these overlaps revealed that genes that have changed expression in only one tissue are rare (∼7%). Instead, most genes have changed in expression across multiple tissues, with the set of genes displaying changes across all five tissues being the largest set by almost twofold (Fig. 2B). Importantly, we find similar results for tissue overlaps and functional category enrichment when identifying differentially expressed genes using a standard alternative (non-phylogenetic) approach, confirming the robustness of our findings (Fig. S4; Methods).

When genes have changed their expression across multiple tissues, this could have occurred simultaneously (e.g., as a result of pleiotropic mutations) or it could have resulted from the accumulation of tissue-specific expression changes at dispersed times in the past (e.g., as a result of the evolution of *cis*-acting regulators or of changes in cell abundances) (Fig. 2C). To gauge the importance of these two contrasting possibilities, we estimated the number of times a gene changed in expression across multiple tissues on individual branches of the phylogeny. Our analysis revealed very few coincidental changes. For example, on the branch leading to *D. melanogaster*, a vast majority of the expression changes occurred in only one of the five tissues (Fig. 2D). The same trend holds when summarizing expression changes over all branches of the phylogeny (Fig. 2E), as well as when quantifying that rate of coincidental changes (Fig. 2F). Collectively, these analyses imply most differentially expressed genes have evolved expression changes across different tissues at independent times in the past, consistent with independent evolutionary changes in gene regulation and/or cellular abundances.

Further inspection of the relatively rare coincidental expression changes indicated that the probability of their occurrence is independent of branch lengths (Fig. S5A). This finding confirms the intuition that many of these expression changes have arisen by pleiotropic mutations (and are not primarily a result of low resolution for detecting independent expression changes along longer branches). Interestingly, the most frequent coincidental change among all tissue combinations involved the forelegs and proboscis+palp samples (Fig. S5B; see also Figs. 2D, E). This observation is coherent with the transcriptomes of these two tissues being the most closely related among the five (Fig. 1B) and points to the likelihood that they share gene regulatory networks.

**Fig. 3.**
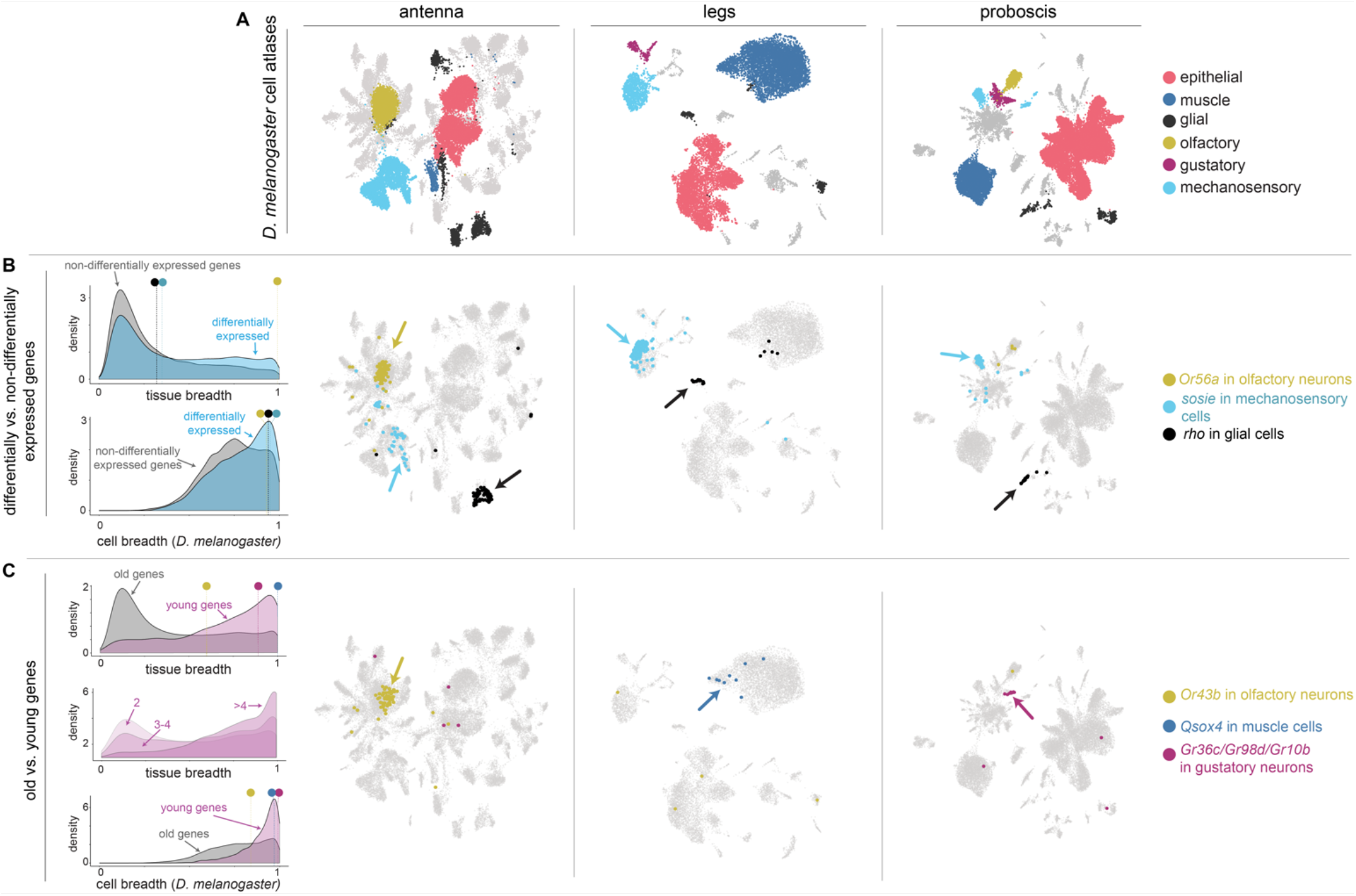
Breadth of gene expression at the level of tissues and cells. **(A)** Single cell atlases from *D. melanogaster* antenna legs and proboscis. Colors highlight the same cell types of interest across tissues. **(B)** Density plots for differentially and non-differentially expressed genes (leftmost panel) relative to expression breadth for tissues (top) and cell types (bottom). Colored circles with lines above the density plots indicate the expression breadth values of three illustrative differentially expressed genes (see text). Expression of the three genes has been mapped onto the *D. elanogaster* cell atlases (right three panels). **(C)** Density plots for old and young genes relative to expression breadth for tissues top left) and cell types (bottom left). The middle left density plot shows the distribution of expression breadth values for genes rouped by duplication levels (2 = paralog group size of 2, 3-4 = paralog group size of 3-4, >4 = paralog group size greater than 4). Colored circles with lines above the top and bottom density plots indicate the expression breadth values of three illustrative new genes (see text). Expression of the three genes has been mapped onto the *D. melanogaster* cell atlases (right three panels).

### Evolution of gene expression is often cell-type specific

Our finding that most differentially expressed genes are broadly expressed led us to question if their breadth of expression differs from genes that have not changed. We first compared the breadth of expression between differentially and non-differentially expressed genes at the tissue level and found that genes that have changed in expression have similar modes of tissue breath but tend to be more tissue-restricted than genes that have not changed (Fig. 3B; Wilcoxon signed-rank test V = 4.3e+10, *p* < 0.001). Based on this result, we hypothesized that differentially expressed genes would also more likely be cell type-specific. Using the recently generated *D. melanogaster* single-cell atlases for antenna, legs, and proboscis^33^, we carried out analogous measures of expression breadth at the level of cell types. Consistent with our hypothesis, differentially expressed genes are indeed significantly more likely to be expressed in a limited number of cell types than genes that have not changed in expression (Fig. 3B).

We found the relationship between tissue breath and cell breath to vary substantially. For example, we identified differentially expressed genes that are expressed narrowly at the cell and tissue levels, e.g., the olfactory receptor *Or56a*, a receptor used by *Drosophilds* to detect the harmful mold odor geosmin^71^ (Fig. 3B). In contrast, we also identified genes that are expressed intermediately at the tissue level but are highly cell-specific within tissues. Using previous cell annotations^33^ and marker-based cell type identification across the three atlases, we verified that these latter cases can be attributed to the same cell types shared across tissues, e.g., *sosie*, a membrane protein localizing to mechanosensory cells and *rho*, a serine protease that localizes to glial cells (Fig. 3B). These examples illustrate how measurements of expression breadth for bulk tissues can mask the cell specificity of a gene’s expression^72^. They also demonstrate that species expression differences that likely underlie phenotypic divergence can be ascribed to individual cell populations.

### New genes tend to be tissue- and cell-specific

New genes are a key source of evolutionary novelty^73^. Due to their potential contributions to species differences, we expanded our analyses to examine how gene age and duplication frequency relate to differences in the transcriptomes of sensory tissues. We compared the breadth of tissue expression between old genes (genes that predated the diversification of the *Drosophila* subgenus more than 50 million years ago^74^) and new genes (genes that arose since^74^). We found that new genes are significantly more likely to be expressed in fewer tissues than old genes (*p*-value < 0.001; Fig. 3C). We also found that the more often a gene has been duplicated, the more tissue-restricted its paralogs are (Fig. 3C). These findings are consistent with the early evolutionary dynamics of new genes^75–79^, and we reasoned that their reduced breadth of expression would correspondingly translate to their detection in a narrower number of cell types. We mapped the expression of new genes to the single-cell atlases for the antenna, legs, and proboscis and compared their breadth of expression across cell types to the expression of old genes. Our analysis confirmed that new genes are indeed significantly more likely to be cell-type specific than old genes (Fig. 3C; Wilcoxon signed-rank test V =3.47e+07, *p* < 0.001).

### Pervasive expression evolution of chemosensory genes

Insect genomes contain three large chemoreceptor gene families: odorant receptors (*Or*s), gustatory receptors (*Gr*s), and ionotropic receptors (*Ir*s)^27^. In addition, members of the chemosensory protein family (*CSP*s), and other diverse protein families, including the odor binding proteins (*Obp*s), transient receptor potential channels (*Trp*s), and pickpocket ion channels (*ppk*s), among others, are chemoreceptors or otherwise involved in the peripheral sensing of environmental chemicals^27, 80–82^. The patterns of expression for most of these “chemosensory genes” have been mapped to specific tissues and cell populations in *D. melanogaster* and have provided the foundation for numerous functional and behavioral studies^27^. While multiple RNA-seq experiments have detected expression differences among developmental stages or species (or both) for chemosensory genes^83–87^, the heterogeneous combination of samples, experimental design, and sequencing approaches have limited evolutionary analyses. We, therefore, manually curated a set of 368 chemoreceptor genes for the above seven gene families and used our uniformly generated RNA-seq dataset to investigate how their expression patterns have evolved between species, tissues, and developmental stages (Table S2).

Out of the 368 chemosensory genes, we detected expression for 299 in at least one of the species’ tissues. In the antenna, proboscis, and forelegs samples, the expression patterns largely matched previous reports^20, 30, 83, 84, 86–114^ (Table S3). However, we detected few of the described *D. melanogaster* chemosensory genes in larva head samples, likely due to their very low expression levels. In each tissue, we identified a core set of genes that were expressed across all six species (antenna = 98, proboscis = 71, forelegs = 63, ovipositor = 28, larva head = 37; Table S4). Fourteen chemosensory genes were found to be expressed across all tissues, including two members of the *ppk* family, *ppk* and *ppk26*, which have previously been implicated in the detection of noxious mechanical stimuli in larvae. Their broad expression suggests additional sensory roles for these proteins in adults. The detection of multiple *Che* members in each of the tissues is also notable, given that their suspected roles in detecting contact pheromones and pathogens have hitherto been limited to the legs^101, 115, 116^.

When we screened the set of 1:1 orthologous chemosensory genes for differential expression, we found that nearly all of them have evolved expression changes in at least one branch of the species tree (Fig. 4A, Table S5; 93% of *CSPs*, 96% of *Gr*s, 100% of *Ir*s, 100% of *Ppk*s, 100% of *Obp*s, 98% of *Or*s, 100% of *Trp*s) Furthermore, most genes have experienced recurrent expression changes, with the *CSP* family showing the greatest. Consistent with genome-wide patterns (Fig. 2A), most expression changes were species-specific. Among these differentially expressed chemoreceptor genes, those that have gained or lost expression in a particular tissue were of particular interest because they may indicate novel gains (or losses) of sensory capabilities. We defined a gene with an average transcript per million (TPM) less than 0.5 as not expressed and a gene with an average TPM greater than 3 as expressed. Using these thresholds, we identified 95 chemosensory orthologs (32%) that have either gained or lost expression in at least one specie’s tissue. Some of these expression gains/losses have occurred once, as illustrated by *Gr98a* and *Gr98b* in *D. melanogaster’s* ovipositor or *D. erecta’s* larva-expressed *Gr59e*, while others have involved recurrent changes, as in the foreleg-expressed *CheB74a* or the antenna-expressed *Ir31a* (Figs. S6-12).

**Fig. 4.**
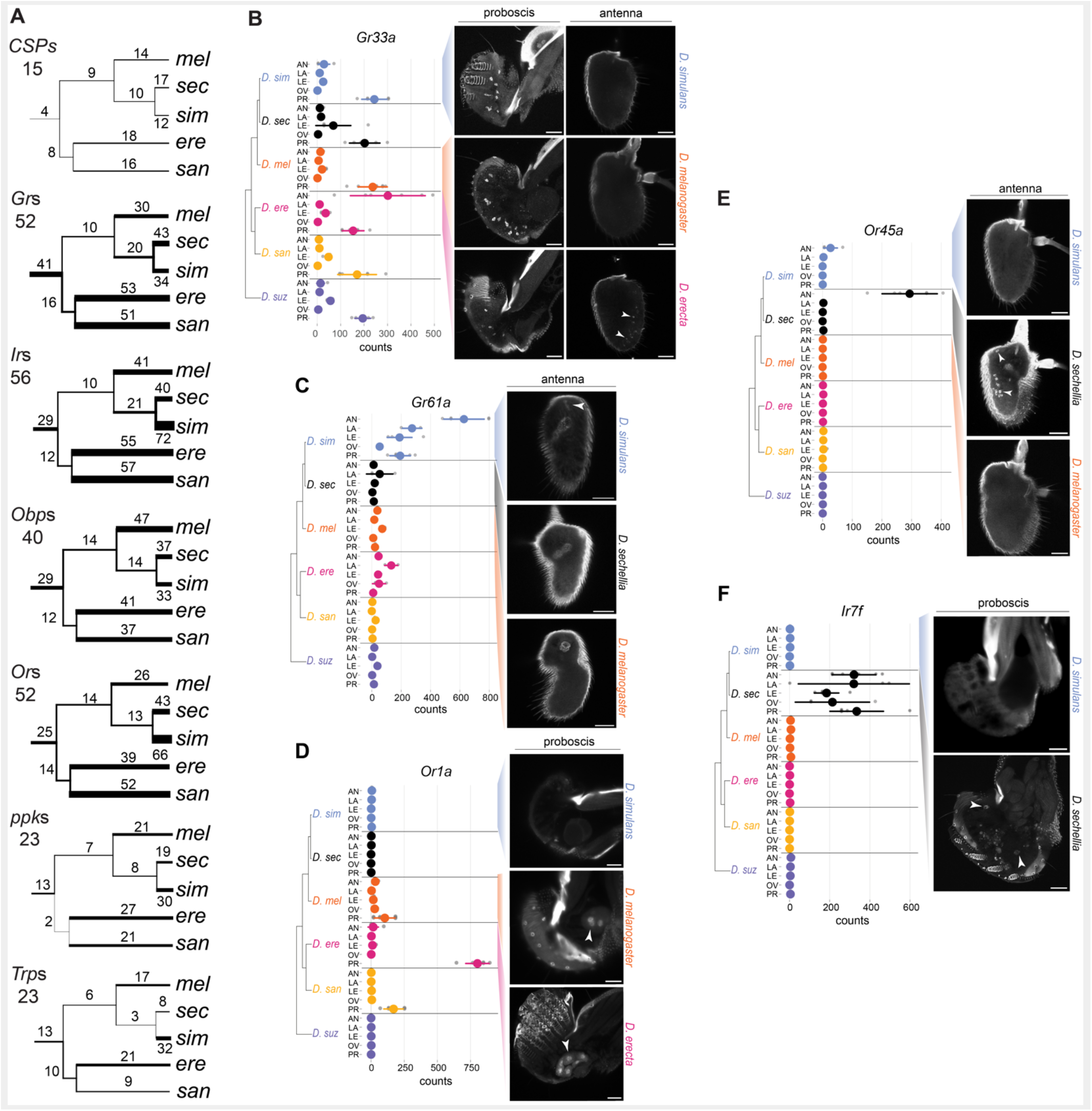
Evolution of chemosensory genes expression. **(A)** Expression changes mapped onto the species tree for genes belonging to the main chemosensory gene families (*Grs* =gustatory receptors, Irs = ionotropic receptors, Ors = odorant receptors, *CSP*s = chemosensory proteins including *CheA* and *CheB* family members, *Obp*s = odorant binding proteins, *ppk* = pickpocket, *Trps* = transient receptor potential channel). The number above each branch is the total number of expression changes (up and down) across tissue samples, with the thickness of the branch proportional to this count. The number under the gene family name corresponds to the number of 1:1 orthologs used for the analysis. **(B-F)** HCR results for chemosensory genes with a species-specific gain of expression. On the left is the species tree (not to scale) with the mean normalized read counts obtained for each sample. The means and standard error are represented by the colored dots and the vertical line, respectively, with the individual data points in grey. AN = antenna, LA = larva head, LE = forelegs, OV = ovipositor, PR = proboscis. RNA *in situ* hybridizations on the right with the targeted tissues(s) above each column. White arrows indicate cells with novel receptor expression. Scale bars: 30µm. See also Fig. S13.

To gain spatial and cellular resolution for the expression of a subset of chemosensory genes with novel expression patterns, we designed *in situ* hybridization chain reaction (HCR) experiments for six of them (*Gr32a*, *Gr33a*, *Gr61a*, *Ir7f*, *Or1a, Or45a*). We detected expression that was consistent with our RNA-seq results for all of these genes except *Gr32a* (Figs. 4B-F). Additional co-labeling experiments resulted in the discovery of several unexpected patterns of cellular expression, including the two gustatory receptors (*Gr32a* and *Gr33a*) expanding their expression to the olfactory system. In *D. melanogaster*, Gr33a was characterized as a bitter receptor expressed in taste cells in the legs and proboscis and involved in aversion to male-male courtship^105, 117, 118^. We found that expression of *Gr33a* has expanded from bitter taste neurons to neurons that express a marker of olfactory neurons (*Orco*) in the antenna of *D. erecta* (Fig. 4B, Fig. S13A). Interestingly, antennal expression of *Gr33a* was previously observed in *D. melanogaster* when programmed cell death was experimentally blocked in olfactory sensory neurons^119^, possibly indicating a *D. erecta*-specific developmental “escape” from cell death for this neuron population. Analogously, *Gr61a*, a glucose receptor in *D. melanogaster* that is expressed in neurons in the labellum, legs, and the labral sense organ^120, 121^, was also found to have expanded into olfactory neurons of *D. simulans*’ antenna (Fig. 4C, Fig. S13B). Analyses of the two odorant receptors (*Or1a* and *Or45a*) also revealed novel expression patterns. We found that *Or1a*, which was previously described as being larva-specific in *D. melanogaster*^107^, is expressed in non-neuronal cells that are likely part of the salivary glands of *D. melanogaster* and *D. santomea* (Fig. 4D, Fig. S13C). To our knowledge, no chemosensory function for this gland has yet been described. We found that *Or45a*, previously described as larva-specific in *D. melanogaster*, is also expressed in the adult antenna in *D. sechellia* (Fig. 4E, Fig. S13D). Finally, *Ir7f*, which has yet to be functionally characterized, was one of the most distinct differently expressed chemosensory genes because it has uniquely gained high expression in all chemosensory tissues in *D. sechellia* (an example of a “coincidental” gain of expression). We observed expression of *Ir7f* within cells that also express a pan-neuronal marker (*nsyb*) in the labial palps, indicating that this gene is likely a taste receptor (Fig. 4F, Fig. S13E). Together, these expression analyses underscore the remarkable evolutionary flexibility in transcript abundance, developmental timing, and spatial expression of chemosensory genes.

### Fast evolution of sex differences

Next, we identified sex differences in our dataset and placed them in a phylogenetic context. *Drosophila* chemosensory tissues are involved in sex-specific functions and often vary between the sexes in morphology and neuroanatomy^19, 69, 122–124^. While previous single gene and transcriptomic analyses identified sex differences in gene expression within some of these tissues^84–86, 125^, their evolutionary histories between tissues and species remain unclear.

For each species, we computed the number of genes with significantly different expression levels between males and females (≥ 1.5-fold change with adjusted p-value < 0.01) within our proboscis+palp, antenna, and foreleg datasets and examined their variation among the six species. Our analysis revealed extensive evolution in the number of sex-biased genes across species, the proportion of genes having male-versus female-biased expression, and in the identity of the sex-biased genes (Fig. 5A; Table S6). Remarkably, the patterns of sex-biased gene expression do not reflect the genetic relationships among the species, in line with previous findings that expression differences between the sexes evolve quickly^126–129^. We observed an approximately ten-fold difference in the number of sex-biased genes between the species with the fewest and the species with the most (*D. santomea* and *D.sechellia* with 135 and 178, respectively, versus *D. erecta* and *D. melanogaster* with 1350 and 1132, respectively). Although the number of male-biased genes outnumbers female-biased genes (2098 vs. 1287), this ratio varied considerably across tissues. Genes expressed in the forelegs and the proboscis are mainly male-biased, while female-biased genes are predominant in antennae. These results suggest that different modes of sexual selection may have shaped the male/female expression balance in a tissue-specific manner.

**Fig. 5.**
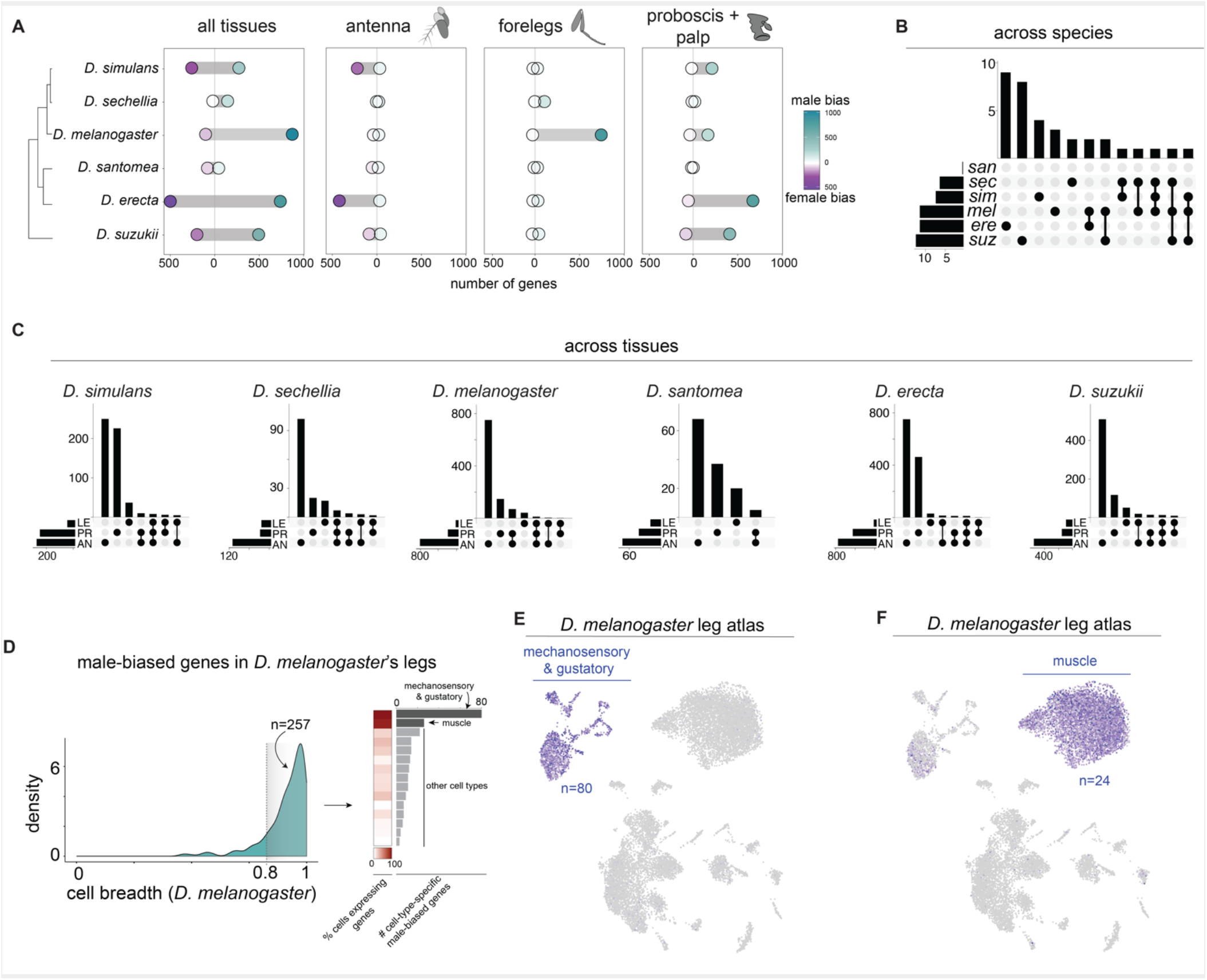
Evolution of sex-biased expression in chemosensory tissues. **(A)** Number of male- and female-biased genes across species and tissues. Sex-biased gene expression does not match the species’ phylogenetic relationships, instead demonstrating many species-specific changes. **(B)** Number of sex-biased genes that overlap across species (regardless of tissue). Most sex-biased genes are species-specific. Species names are abbreviated to their first three letters. **(C)** Number of sex-biased genes that overlap across tissues within species. Most sex-biased genes are tissue-specific (LE=forelegs, AN=antenna, PR=proboscis+palps). **(D)** (right) Density plot for *D. melanogaster’s* male-biased genes relative to cell-type expression breadth. (left) heat map showing the fraction of cells in a given cell population that express the male-biased genes and bar plot displaying the total number of male-biased genes found expressed within a given cell population. Most male-biased genes are cell type-specific and predominantly found within cells associated with mechanosensation, gustation, and muscles. **(E)** Cell atlas for *D. melanogaster* legs showing the total mean expression of 87 male-biased genes. Their expression is restricted to mechanosensory/gustatory cells. **(F)** Cell atlas for *D. melanogaster* legs showing the total mean expression of 24 male-biased genes. Their expression is restricted to muscle cells.

We then asked if the identity of sex-biased genes is shared across species and tissues. These analyses once again highlighted pervasive variation in the sets of genes that differ between the sexes. In a majority of cases, the sets of sex-biased genes are private to each species (Fig. 5B). Intriguingly, among the few overlaps, we found enrichments of genes involved in or activated by cell-autonomous and non-autonomous control of sex differences (including *fruitless*, *doublesex*, *insulin-like peptide 7*, and members of the cytochrome P450 family; hypergeometric tests *p*-values < 0.001), suggesting that they may play roles in the maintenance of sexually dimorphic traits in some adult sensory tissues, similar to what has been observed for *D. melanogaster’s* intestine^21^. We also observed enrichment in chemosensory proteins (hypergeometric tests *p*-values < 0.001) which have been shown to be sex-biased and involved in pheromone-induced behaviors^101, 115^. Finally, within species, if a gene is sex-biased in one tissue, it is rarely sex-biased in the other two (Fig. 5C).

We sought further insight into the cell types underlying the derived *D. melanogaster* male-biased foreleg expression. Of the 806 male-biased genes, 285 were present in the leg cell atlas^33^. Plotting their distribution in relation to cellular expression breadth revealed remarkable cell-type-specificity with the most cell-type-specific genes (cell breath >0.8; n=257) enriched in cell populations related to mechanosensation, gustation, and muscle (Fig. 5D-F). These cell types are particularly compelling in light of the extensive literature identifying key roles for these sensory modalities in *D. melanogaster*’s courtship^22, 130–132^, and because musculature is dimorphic among *Drosophilids*^133^. Genes involved in mitochondrial respiratory (*ND-88*) and actin assembly (*forked*) are found with the muscle population, as are two vision-related genes (*Rh2* and *Culd*). Among characterized genes in the mechanosensation/gustation cell population are a putative pheromone receptor (*Ir52c*), two *trp* channels involved in temperature sensing (*brv3* and *pdk2*), and several genes involved in neuron development and signaling (e.g., *Unc-104* and *Stathmin*). As the Fly Cell Atlas contains a male and a female sample (two samples each and pooled), we next asked whether cellular composition or transcript abundance differs between the sexes. Though preliminary, we found more muscle cells that are enriched for the male-biased genes in the male sample compared to the female sample and also found higher mean expression levels for the male-biased genes in the male sample (Wilcoxon test *p <* 0.01). We did not find differences between sexes in the mechanosensation/gustation cell population (Fig. S14). These results suggest that, at least in the muscle cells, regulation of the cell population size and transcript abundance have contributed to this sexual dimorphism in *D. melanogaster*.

## Discussion

By conducting comparative transcriptomic analyses of chemosensory tissues across species, and linking them with single-cell datasets and additional experiments, we have expanded our understanding of how these sensory systems evolve on a global and individual gene level. Globally, we have found that the chemosensory transcriptomes are shaped predominantly by stabilizing selection. Nevertheless, the broad evolutionary constraint has not precluded a subset of tissues and genes from experiencing accelerated rates of expression change. At the transcriptomic level, *D. melanogaster* (forelegs and ovipositor) and *D. simulans* (antenna, larva head, and ovipositor) are distinct for having significantly increased expression divergence. This is initially curious as the two are ecological generalists while the other species have evolved ecological specializations. However, it is consistent with *D. melanogaster* and *D. simulans* having the largest effective population sizes (and likely substantially so)^47, 52^ resulting in positive selection playing a greater role within these species compared to the others. If true, this result would suggest an important role for positive selection in driving gene expression changes. At the level of individual genes, we have identified numerous expression changes across the chemosensory tissues of all six species. Most of these changes have occurred in only one species, indicating that many of the expression differences are recent. As it becomes more feasible to carry out population surveys for expression polymorphism, it will be important to quantify how many of these changes are fixed between species and how many are polymorphic^129, 134^.

The expression changes that we identified could have resulted from species’ differences in transcript abundance (e.g., *cis*-regulatory changes) or cellular composition (e.g., expanded or contracted cell populations). Though we cannot separate the possibilities with bulk tissue samples, the fact most of the changes occurred tissue-specifically (“dispersed”) supports an evolutionary model of modular change. This observation is important because a key factor in determining anatomical evolution is the pleiotropy of mutations. Due to the functions that individual genes have across multiple tissues, it is expected that the diversification in any one tissue (or subset) will arise through mutations in genes’ modular cis-regulatory regions^67, 135^. To the extent that transcript abundance drives the differences in our datasets, our results are consistent with previous findings that indicate that most between-species expression changes are driven by *cis*-regulatory modifications^136–140^. We expect that the close relationships between these species will foster the identification of candidate regulatory differences that can be studied to further understand the molecular basis of transcript abundance evolution. Much less is known about the evolution of cell population sizes. In the case of *D. melanogaster*’s novel male-biased foreleg expression, we have found preliminary evidence that both transcript abundance and cell composition evolution may be involved. We will soon be able to address this question more thoroughly through cross-species comparisons of single-cell atlases.

Molecular evolutionary studies of chemosensory genes have consistently highlighted their rapid protein coding and copy number evolution^141–144^. Our analyses demonstrate that changes in transcript abundance and novel expression gains and losses also fuel their fast evolution. It has been suggested that the cell-specific expression patterns of most chemosensory genes, along with partially overlapping molecular functions (e.g., promiscuous ligand-binding), result in relatively fewer pleiotropic constraints and, as a result, increased evolutionary freedom to change^141^. It is likely that their narrow cellular expression also allows for increased flexibility to fine-tune their levels of expression. Though the phenotypic implications of chemoreceptor expression levels remain unclear, it is plausible that they shape neuronal sensitivity or other cellular kinetics that impact a neuron’s encoding of chemical information. We also have evidence for several peripheral sensory neurons population that they can expand/contract quickly^54, 145^ and are likely contributing to species differences in chemoreceptor expression levels. More comprehensive studies are needed to assess how frequently such changes are occurring. Of potentially greater immediate phenotypic consequences are chemoreceptors’ ability to gain (or lose) expression in novel tissues. We estimated that approximately a third of the chemosensory genes may have done so over the diversification of these six species. And while instances of unusual or “ectopic” receptor expression, as illustrated by *Or1a* (Fig. 4D), call for additional functional characterization, they are also a reminder of the first step that all receptors and receptor operated channels have taken as they have diversified across tissues throughout the animal kingdom.

As with other comparative functional genomic studies, identifying the specific changes that are translated into phenotypic differences remains an outstanding challenge. The phylogenetic framework provided here will help to devise future experiments for addressing this question, as illustrated by our investigation of five chemosensory genes with novel expression patterns. One line of evidence pointing towards a substantial fraction of the expression differences being functionally important is our observation that they tend to be cell-specific. Though it is conceivable that a similar trend could be produced by neutral evolution (e.g., expression drift being more common among sets of genes that are cell-type-specific), we argue that this observation nonetheless provides new and important genome-wide evidence consistent with them being functionally relevant. This is most convincing in the context of sex differences, where nearly all expression changes are species-specific and where, in *D. melanogaster*, we linked species-specific sex-biased genes to specific cell populations involved in sexually dimorphic functions^22, 130–132^.

## Methods

### Fly strains, rearing, and dissections

Fly strains (*D. melanogaster* B54^146^, *D. simulans* 14021-0251.008, *D. sechellia* 14021-0271.07, *D. santomea* 14021-0271.00, *D. erecta* 14021-0224.01, *D. suzukii* K-AWA036) were reared on a standard yeast/cornmeal/agar medium supplemented with Carolina 4-24 Formula and maintained in a 12:12 hr light:dark cycle at 25 degrees. Adults between 2 to 10 days old were sex-sorted on CO2 at least 24h before the dissections. Third instar larvae were taken directly from the food medium the day they were dissected. For each replicate, 10 third instar larval heads, 25 proboscis, 50 legs, 5 ovipositors, and ∼100 antennae were collected. Three replicates were made per sex and species for the proboscis, the legs, and the antennae; 3 replicates were made per species for ovipositors and larval heads.

### Tissue collection

All adult samples were collected from flies aged between 2-10 days. Antennas were collected by flash-freezing flies in liquid nitrogen and agitating them over a mini-sieve connected to a collection dish^147^. Antennas were selected from the collection dish using a pipette under a dissecting scope. Forelegs and proboscis with maxillary palps were collected from individual files using forceps and a micro scalpel under a dissecting scope. Third instar larvae were collected from vials by floating them in 75% sucrose water and washed. Larva heads were removed under a dissecting scope using a micro scalpel.

### mRNA library preparation and sequencing

Dissected tissues were homogenized in 200ul of Trizol (Invitrogen) using a Precellys24 (6800rpm, 2×30s with 10s breaks; Bertin Technology) followed by a standard Trizol RNA extraction. The final mRNA concentration was measured using a DeNovix Ds-11 FX spectrophotometer. mRNA libraries were prepared using KAPA Stranded mRNA-seq Kit (Roche) following the manufacturer’s instructions (Version 5.17). Briefly, 500ng of total RNA diluted in 50ul of RNAse-free water was first placed on supplied mRNA capture magnetic beads to allow the isolation of mature, polyadenylated mRNA, which was subsequently fragmented to a size of 100-200bp. Double-strand cDNA was then synthesized, marked by A-tailing and barcoded with 2.5ul of TruSeq RNA UD Indexes (Illumina). SPRI select beads (Beckman Coulter) were used for cleanup. Library concentrations were measured using Qubit dsDNA HS Assay Kits (Invitrogen). Fragment analysis and HiSeq 4000 single-end Illumina sequencing were performed by the Lausanne Genomic Technologies Facility.

### *In situ* hybridization chain reaction experiments

*Probe sets:* HCR probes set, amplifiers, and buffers were purchased from Molecular Instruments. The list and the sequences of the probes used can be found in Table S7. Coding sequences and 5’ and 3’UTRs, were extracted from the species reference genomes and aligned. *D. melanogaster* sequences were used to design HCR probe sets for genes sharing >91% identity across our target species. If sequence identity was less than 91%, or if we failed to detect a signal using a *D. melanogaster* probe set in a different species where transcripts were detected in our RNA-seq dataset, we designed species-specific probe sets. Based on these criteria, species-specific probes were designed for *D. simulans Gr61a*, *D. suzukii Gr32a*, *D. sechellia Ir7f,* and *D. suzukii Gr66a*.

*In situs*: Flies between 2 to 9 days old were cold anesthetized and dissected on ice. Samples were collected on PBT (1XPBS, 0,1% Triton X-100) and fixed for 2h or 24h in 2ml of 4% paraformaldehyde, 1X PBS, 0.1% Triton X-100 at 4 °C on a rotator set at low speed (<20 rpm). Following fixation, samples were washed twice in PBS + 3 % Triton X-100 and three times in PBT. The protocol suggested by Molecular instruments for generic samples in solution was then followed with minor adjustments (https://files.molecularinstruments.com/MI-Protocol-RNAFISH-GenericSolution-Rev9.pdf). Samples were pre-hybridized in 300μl of probe hybridization buffer for 30min at 37°C. For antenna samples 3,5μl of control probe (*Orco* or *nsyb*) and 5μl of experimental probes were used. For proboscis, 5ul of control (*nsyb*, *Gr66a*) and experimental probes were added to the amplification buffer. Samples were also pre-amplified in 300μl of amplification buffer. For antenna samples 6μl of hairpin solution designed to amplify the signal of control probes was used, 10μl otherwise. For proboscis samples, 10μl of hairpin solutions were used to amplify both the controls and the experimental probes. After washes, samples were mounted in Vectashield and stored at 4°C.

*Image acquisition:* Antennae, proboscis and larvae images were acquired on inverted confocal microscopes (Zeiss LSM 710 or LSM 880) equipped with an oil immersion 40× objective (Plan Neofluar 40X oil immersion DIC objective; 1.3 NA). The images were processed in Fiji (v1.53)^148^.

### Gene annotations

Annotations in General Feature Format were generated for all species using BRAKER v2.1.6 and Augustus v3.4.0^149, 150^. We ran BRAKER with the --etpmode flag as we provided evidence from both our aligned RNA-seq data and an orthologous protein dataset for arthropods (arthropoda_odb10). The quality of annotations was checked with BUSCO v3.0.2^151^. First, we generated fasta files with coding sequence from the annotations using Cufflinks v2.2.1^152^ gffread function (`-w exons.fa -W -F -D -E -o filtered.gff flags`). Completeness was checked against the diptera_odb9 dataset. BUSCO scores were similar across species: *D. simulans* 97.3%, *D. melanogaster* 97.1%, *D. erecta* 97.0%, *D. santomea* 94.5%, *D. suzukii* 91.9%. The species’ GTFs are in File S1.

### OrthoFinder-based orthology analysis

Our next goal was to group our annotated sequences into their respective orthologue groups using OrthoFinder v2.3.8^153^. The input peptide sequence was generated for each species by the following steps: (1) fasta files of coding sequence from annotations were converted to peptide sequence using the transeq function from EMBOSS v6.6.0^154^, (2) duplicate genes introduced from BRAKER’s pipeline were removed using a custom script (rmduplicategenes.sh), (3) Orthofinder’s primarytranscript.py was run on each of the resulting peptide fasta files. These input peptide sequences were then placed in the same directory and we ran OrthoFinder to generate our orthologue groupings. We additionally added the -M msa flag to generate gene trees

### Opposvum-based orthology analysis and gene IDs

We used Possvm^155^ (v1.1) to refine orthology relationships (1 to many and many to many) inferred by Orthofinder (above). We first aligned non-1:1s orthologs using MAFFT^156^ (v7.490; `mafft --auto protein.fa`) and outputted alignments in phylip format. We then generated phylogenetic trees containing bootstrap information at each node using IQ-TREE^157^ (v2.2.0.5; `iqtree2 –s ./MAFFT_ortho/${spe} -mset WAG,LG -b 200`), testing for the best substitution model (WAG or LG) and performing 200 bootstrap replicates. Finally, we used Possvm to identify new orthogroups. For this step, we first parsed phylogenies using the species overlap algorithm, and second, we clustered orthogroups using the MCL clustering method. We updated the former orthofinder Orthogroup.tsv with the list of newly generated orthogroups which included 2,066 new 1:1s genes. Orthogroups were renamed according to the *D. melanogaster* reference genes, which were identified through iterative BLAST (v2.10.1+)^158^. For this step, we used tblastn to query our list of protein orthogroups on a *D. melanogaster* gene database containing nucleotide fasta from all annotated CDS. BLAST results were sorted according to their best hit (bit score selection) and the matching gene names were appended to our inferred orthogroups IDs. This “lookup table” is available as Table S8.

### RNA-seq read mapping

Each species’ Illumina reads were mapped to its own soft-masked reference genome using STAR (v2.7.8^159^), inputting the GTF files generated above. On average we obtained 43 million mapped reads per replicate, with mapping rate modes ranging between 0.79-0.90. A single *D. simulans* proboscis sample resulted in fewer mapped reads due to the amplification of a viral sequence, but was otherwise highly correlated with the two other replicates and was therefore retained. Sample replication across all tissues was high, with an overall average Pearson correlation coefficient of 0.98; the range of Pearson correlation coefficients within each tissue’s replicates was between 0.97-0.99. The one exception was the ovipositor dataset, likely reflecting less precise dissections (above). Pearson correlation coefficients for the ovipositor samples ranged from 0.93-0.99, with the replicates of *D. erecta*, *D. santomea*, *D. sechellia* being more variable (average Pearson correlation coefficients = 0.93, 0.95, 0.96, respectively) than the other three species (average Pearson correlation coefficients = 0.97, 0.97, 0.99).

### Read count table generation

*Full-length gene*: Expression count tables were generated using HTseq (v0.11.2^160^), inputting the GTF files generated above (File S2; the corresponding TPM table for the 1:1 orthologs is File S3).

*Trimmed genes:* Despite the six species being closely related, differences in orthologous gene lengths exist. If unaccounted for, these differences may lead to misleading cross-species differential expression results when using methods that assume identical gene lengths. To account for length differences in our PCA or clustering analyses and for analysis using DESeq2 (v1.34.0^161^), we generated count tables based on orthologous gene regions that were conserved across all six species. Conserved regions were identified based on DNA alignments (MAFFT v7.475^156^) of the 1:1 orthologs. We excluded gene regions if any of the six species contained a gap greater than 150bp (using the script get_aligned_blocks.py). Using the coordinates of the conserved gene regions, we then generated a set of “trimmed” GTF files (using the script make_trimmed_gtf.py; the species’ trimmed GTFs are found in File S4) that were passed to HTseq (v0.11.2^160^) for computing the “trimmed” count tables (File S5). The “trimmed” GTF file includes the full set of genes that were annotated in each species’ genome but contains the modified genic coordinates based on the conserved alignments for the set of 1:1 orthologs.

### Transcriptomic clustering and Relative rate tests

Transcriptomes were clustered by species using the set of 1:1 orthologs and a phylogenetically-informed distance measure implemented in TreeExp (v0.99.3^70^). TreeExp implements a statistical framework assuming that gene expression changes are constrained by stabilizing selection (based on the Ornstein-Uhlenbeck (OU) model). For phylogenetic reconstruction, we generated “taxa.objects” from our TPM normalized expression matrix specifying taxa (species) and sub-taxa (tissue) levels. Distance matrices were computed for each tissue by modeling gene expression changes under a stationary OU model (method= “sou”). Finally, distance matrices were converted into phylogenetic trees using the neighbor-joining method, setting *D. suzukii* as an outgroup and performing 100 bootstrap replicates.

Relative rate tests were carried out in TreeExp (v0.99.3^162^) for all pairwise comparisons using its RelaRate.test function. For these analyses, only genes with a TPM >1 were included. To confirm that divergence score estimations were not driven by a subset of genes as well as to give stronger statistical power to the analysis, we computed divergence Z-scores by randomly sampling 1,000 genes and 1,000 times for each species pair and each tissue sample. We compared the per species per tissue Z-score distribution from randomly sampled genes to both the minimum and maximum value of the non-significant Z-score distribution using a Wilcox test statistic in R^163^.

### Differential expression for 1:1 orthologs

*OU analyses*: Evolutionary changes in gene expression were detected using the l1ou R package (v1.43^164^). The method uses a phylogenetic lasso method to detect past changes in the expected mean trait value, assuming traits evolve under the OU process. We used a reference species tree that was previously inferred^53^ and the species’ mean TPM for each gene, for each tissue, as the evaluated traits. We set the maximum number of possible expression changes to 3 (half the number of taxa in the tree) and selected the best model for the number of expression changes using the phylogenetic-informed BIC approach (pBIC).

*Coincidental index:* For each gene, we calculated the frequency that it changed in expression in multiple tissues simultaneously by computing a simple “coincidental index” defined as:

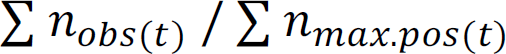

Where *n_obs(t)_* is the number of tissues an expression change occurred at time *t* and *n_max.pos(t)_* the maximum number of possible changes at time *t*. This index takes a value between 0 to 1 where 0 reflects no expression changes, 0.2 reflects a change that occurred in only one tissue (dispersed) and 1 an expression change that occurred simultaneously in all tissues (coincidental).

*DESeq2 analyses*: Pair-wise based identification of differentially expressed genes was carried out with DESeq2 (v1.34.0^161^) specifying the following design: ∼ 1 + species + tissue + species:tissue. For these tests, the set of 1:1 orthologs (above) and the “trimmed” count tables (above) were inputted.

*Gene module analyses:* We identified co-expressed gene modules between tissues using soft-clustering algorithms implemented in CEMITools^165^.

### Analyses of gene duplicates and expression

We performed gene age analyses on gene lists derived from^166^. Genes predating the speciation of the *Drosophila* subgenus (∼50 My ago) were classified as “old”, while new genes that have emerged since the *Drosophila* subgenus speciation event were classified as “young”. Duplicated genes and their level of duplication are derived from our ortholog annotation on the set of non-1:1 orthologs (Table S8).

### Manual curation of chemosensory gene set

Chemosensory genes were first extracted from the look-up table generated for the global dataset (Table S8). Genes for which an ortholog was missing in one or more species, genes with multiple paralogs, or genes previously annotated in *D. melanogaster* or *D. suzukii* but missing in our datasets were investigated and manually corrected if an annotation error was identified. *D. melanogaster* coding sequences were obtained from flybase^167^ and *D. suzukii* coding sequences from the literature^143^. Each species’ reference genome was uploaded into Geneious (v2022.0.2) and annotated with the GTF files generated above. The coding sequences of the genes selected for manual correction were then combined with these annotated genomes using Minimap2 (v2.17^168^). A new GTF file for the chemosensory genes was generated for each species with annotation errors corrected and previously omitted missing genes added. The GTF files for these manually curated annotations are available in File S6.

For each tissue, the mean TPMs for each gene across replicates were calculated and the number of genes from each chemosensory family that were detected as expressed was evaluated with TPM thresholds of 0.25, 0.5, 1, 2, and 3. For the antenna and proboscis, the number of genes detected only slightly decreased with TPM thresholds between 0.5 and 2 TPM. However, for ovipositor, forelegs, and larva, the number of genes detected dropped significantly with the increase of the TPM threshold. This is likely because some genes are expressed in a few cells, leading to low TPM values. Therefore, to ensure that these genes were not excluded, the threshold for gene detection was settled at 0.5 TPM for all tissues. The TPM file for the chemosensory set of genes is available in File S7.

### Sex-biased gene expression

Genes that have significant differences between sexes were identified using the full set of species’ genes and the “full gene” count tables. Read count data for the tissues of each species was read into DESeq2 (v1.34.0^161^) specifying the following design: ∼ tissue + sex + tissue:sex. Only genes that had a normalized read count of five in three or more samples were kept for analysis. A Wald test was used to test for sex differences for each gene, requiring a log fold change of 1.5 and an adjusted *p*-value < 0.01 for significance.

### Fly Cell atlas data manipulation

*Data importation*: We imported 10x stringent loom and H5DA atlases of legs proboscis and antennas from flycellatlas.org^33^. The H5DA files (that contain the clustering information and feature count matrix for a subset of Highly Variable Genes) were converted to Seurat objects using the Convert function from the SeuratDisk (v0.0.0.9020; https://mojaveazure.github.io/seurat-disk/) and were exported as RDS files using the “saveRDS” function. We used the “Connect” function from SeuratDisk to convert loom files (containing count matrix for all *D. melanogaster* genes but no clustering information) to Seurat objects and exported them as RDS files.

*Mean gene expression per cell cluster*: We split Seurat objects by cluster (`subset(atlas.data, idents=“cluster_ID”)`) and extracted their respective feature count matrices (`GetAssayData(object=atlas.data, slot=“count”)`). We then calculated the mean expression of individual features per cluster (`rowMeans()`) and log-transformed their expression for downstream analysis.

*Visualization of a gene of interest*: The AddModuleScore function from Seurat was used to select gene subsets and visualize their expression using the “FeaturePlot” function. To visualize subsets of cells expressing specific features, we used the “DimPlot” function specifying cells of interest with the “cells.highlight” option. Gene expression cutoffs were determined after visual examination to highlight highly expressing cells only.

*Cell type homology between tissue*: We used the Seurat “FindAllMarkers” (`atlas.data, only.pos = TRUE, min.pct = 0.25, logfc.threshold = 0.25`) function from Seurat to identify significant markers (*p*-value < 0.001) among the top 100 list of markers per cell cluster. The list of unique shared markers was retrieved across all tissues, and we generated pairwise correlation matrices based on cluster-mean expression values for each cell cluster across each tissue. In addition, we generated pairwise matrices of the percentage of cell markers shared across tissue cell clusters. The product of these two matrices gives a score between 0 and 1, where 0 corresponds to completely unrelated cell types, and 1 corresponds to identical cell types. This homology score enabled us to cross-validate the FlyCellAtlas annotation and to identify cell type homology across tissues at a finer scale.

### Measurements of tissue breadth and cell type breadth

We measure gene expression breadth using the summary statistic τ^169^ defined as:

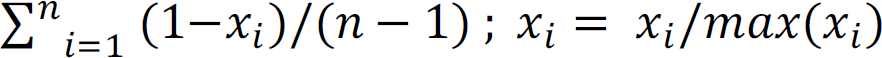

Where *x_i_* is the expression of the gene in tissue *i*, *n* the number of tissues.

We apply the same formula to define τ at the level of cell clusters where *x_i_* is the mean expression of the gene in cell cluster *i* and n the number of clusters in a given atlas. We also investigated measuring cell specificity τ index by considering *x_i_* as the percentage of cells expressing the gene in cluster *i,* which gave very similar distributions. All count values were log-transformed before applying the τ formula for stringency purposes.

### Reagents, code, and data availability

Information for all molecular reagents used in this project can be found in Table S7. Beyond supplementary data, all code and data used in this project are available on our lab’s “sensory RNAseq” GitLab repository: https://gitlab.com/EvoNeuro/sensory-rnaseq. All fastq files generated for this project are available on ArrayExrpress under the accession code E-MTAB-12656. The normalized count data for 1:1 orthologs can be explored and plotted with our CT^2^ dashboard available at: https://evoneuro.shinyapps.io/ctct/.

## Acknowledgements

We thank Margarida Cardoso-Moreira for discussions and advice throughout the completion of the project. Margarida Cardoso-Moreira, John Pannell, Thomas Auer, Giulia Zancolli, and Lucia L. Prieto Godino provided important comments on an earlier version of the paper. Sequencing was performed at the Lausanne Genomic Technologies Facility, and the University of Lausanne’s HPC services provided resources and support. Research in JRA’s laboratory was supported by the University of Lausanne and the Swiss National Science Foundation (grants PP00P3_176956 and 310030_201188).

## Author contributions

Conceptualization: GB, BSL, JRA

Investigation: GB, BSL, TK, TB, AH, JASA, JRA Visualization: GB, BSL, JRA

Funding acquisition: JRA Supervision: JRA

Writing – original draft: JRA

Writing – review & editing: GB, BSL, TK, TB, AH, JASA, JRA

## Supplemental Figures

**Fig. S1.**
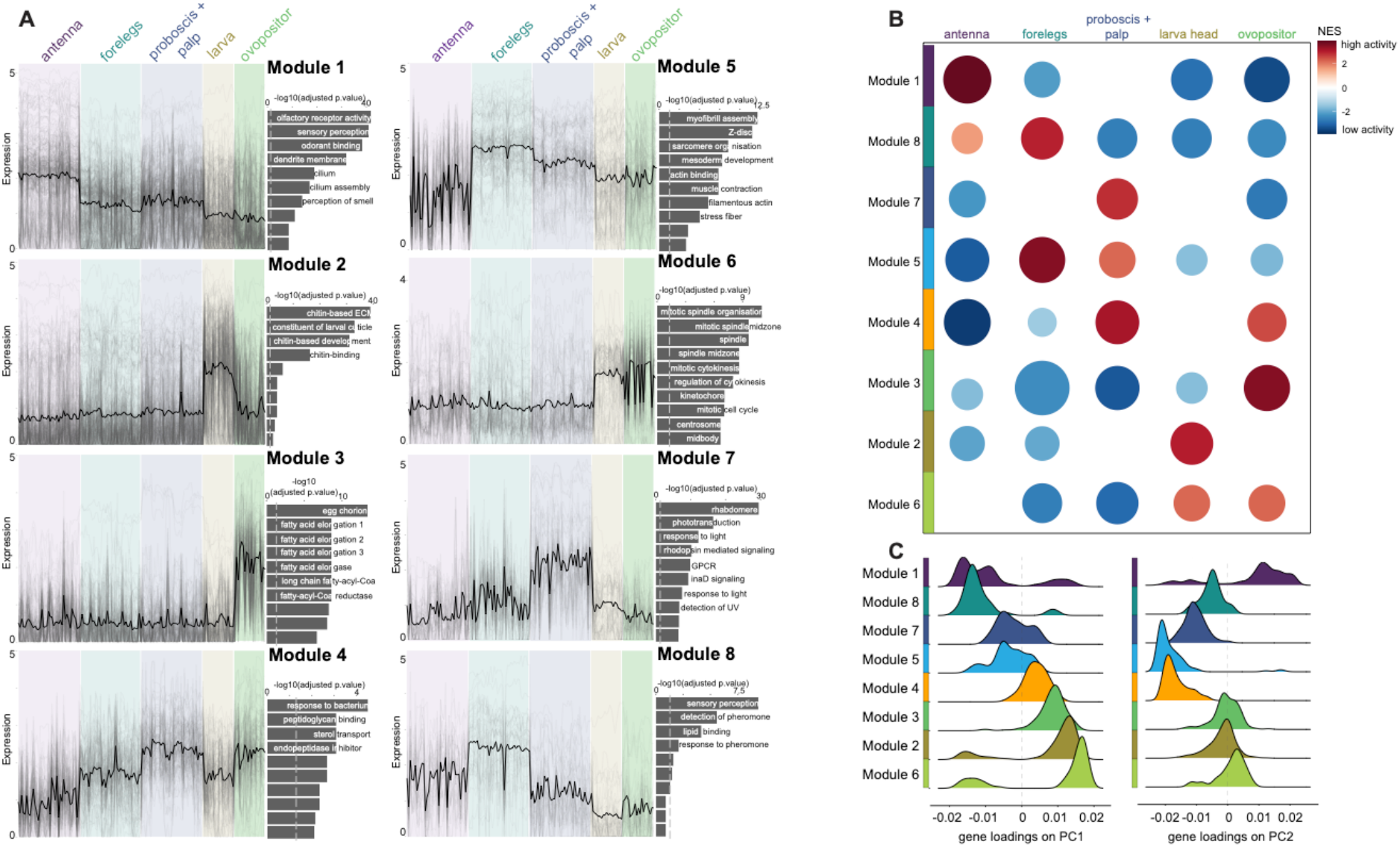
Identification of expression modules across chemosensory tissues. We used CEMITool to identify co-expressed gene modules. From gene expression table, CEMITool uses unsupervised filtering method to select genes used in the analyses. It then uses soft-clustering methods to determine a similarity criterion between pairs of genes. Based on this criterion, genes are separated into modules unsing the Dynamic Tree Cut package. **(A)** Gene co-expression analyses showing expression profile of individual genes (thin lines) across samples group by tissue (left plots). The thick line displays the median expression of all co-expressing genes within a gene module. Right to profile plots are histograms of enriched pathways ranked by *p*-values. Dashed lined show significance thresholds. **(B)** Get Set Enrichment Analyses displaying the modules’ (from panel A) activity per tissue. **(C)** Density plots showing the distribution of genes belonging to a module (from panel A) on the first (left) and second (right) principal components of the PCA from Fig. 1B.

**Fig. S2.**
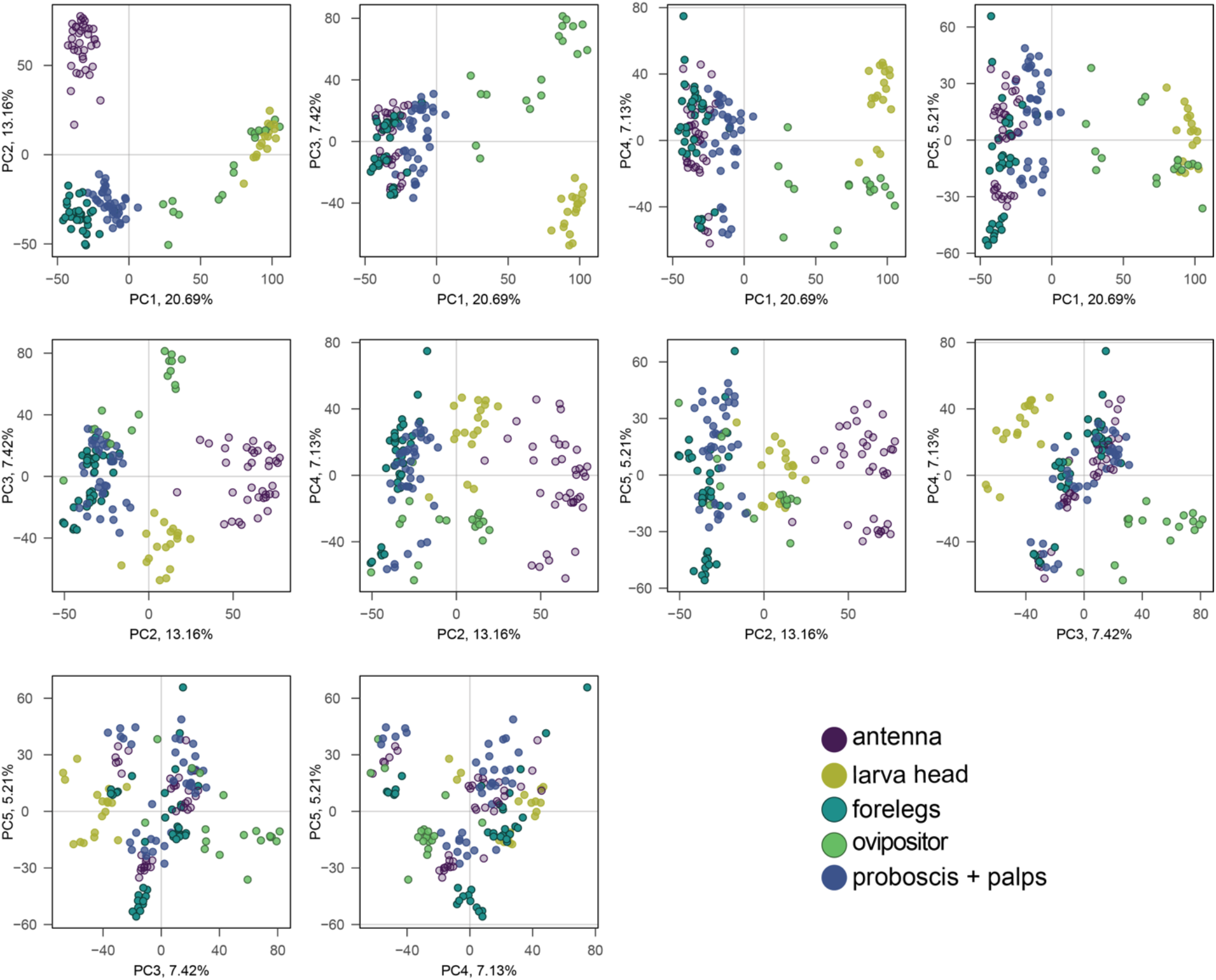
PCA analysis of chemosensory transcriptomes beyond PC 1 and PC 2. Across the different principal component pairings, the only two tissues that do not separate are the foreleg and proboscis+palps.

**Fig. S3.**
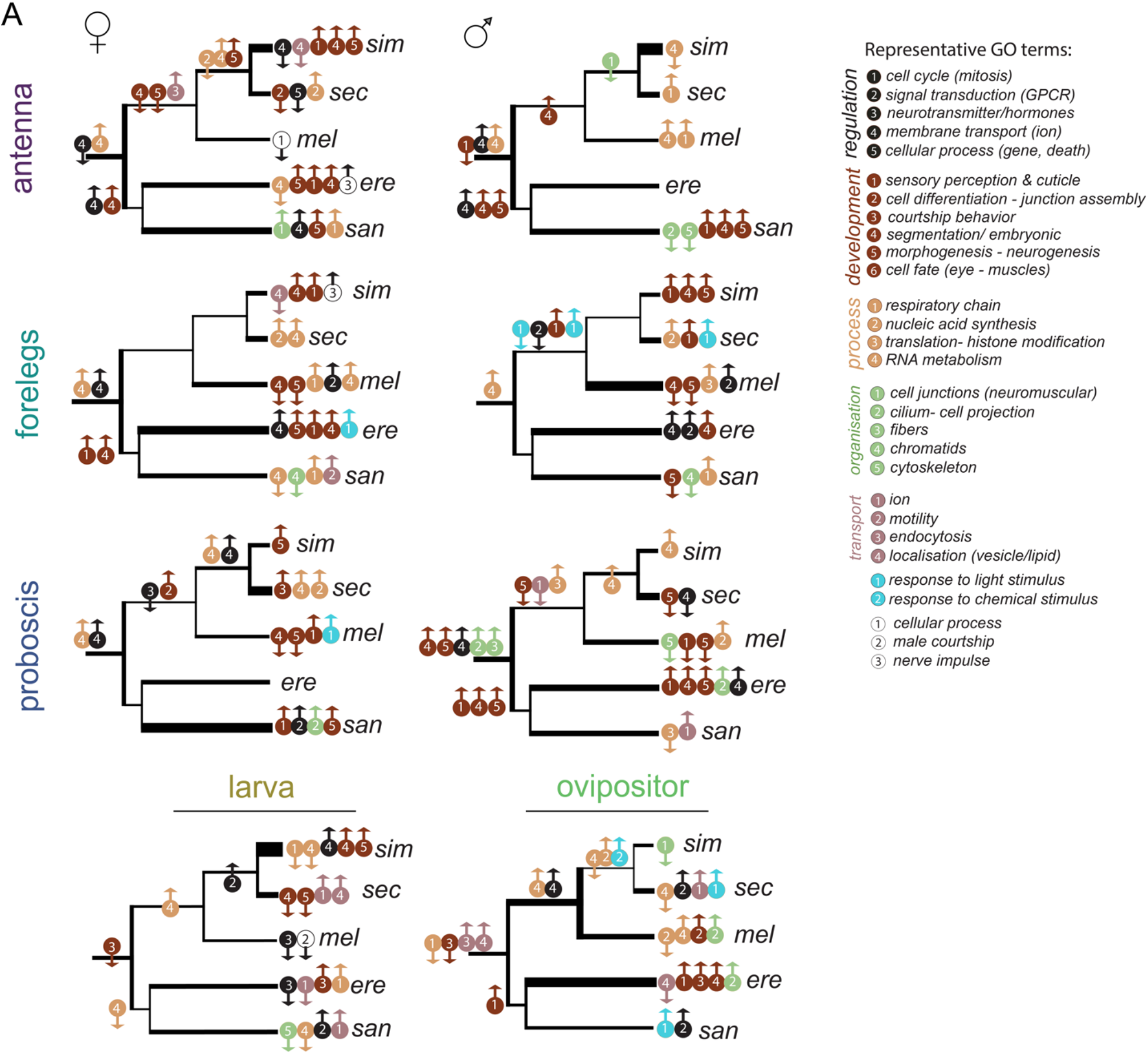
Functional enrichment analysis of differentially expressed genes. **(A)** GO term enrichment analyses of differentially expressed (DE) genes for each branch of the species tree and each tissue. Circles correspond to representative terms listed on the right panel. Circles are colored according to large semantic categories and numbers correspond to semantic sub-clusters within these categories. We performed GO term analyses for up-regulated (up arrows) and down-regulated (down arrows) genes separately.

**Fig. S4.**
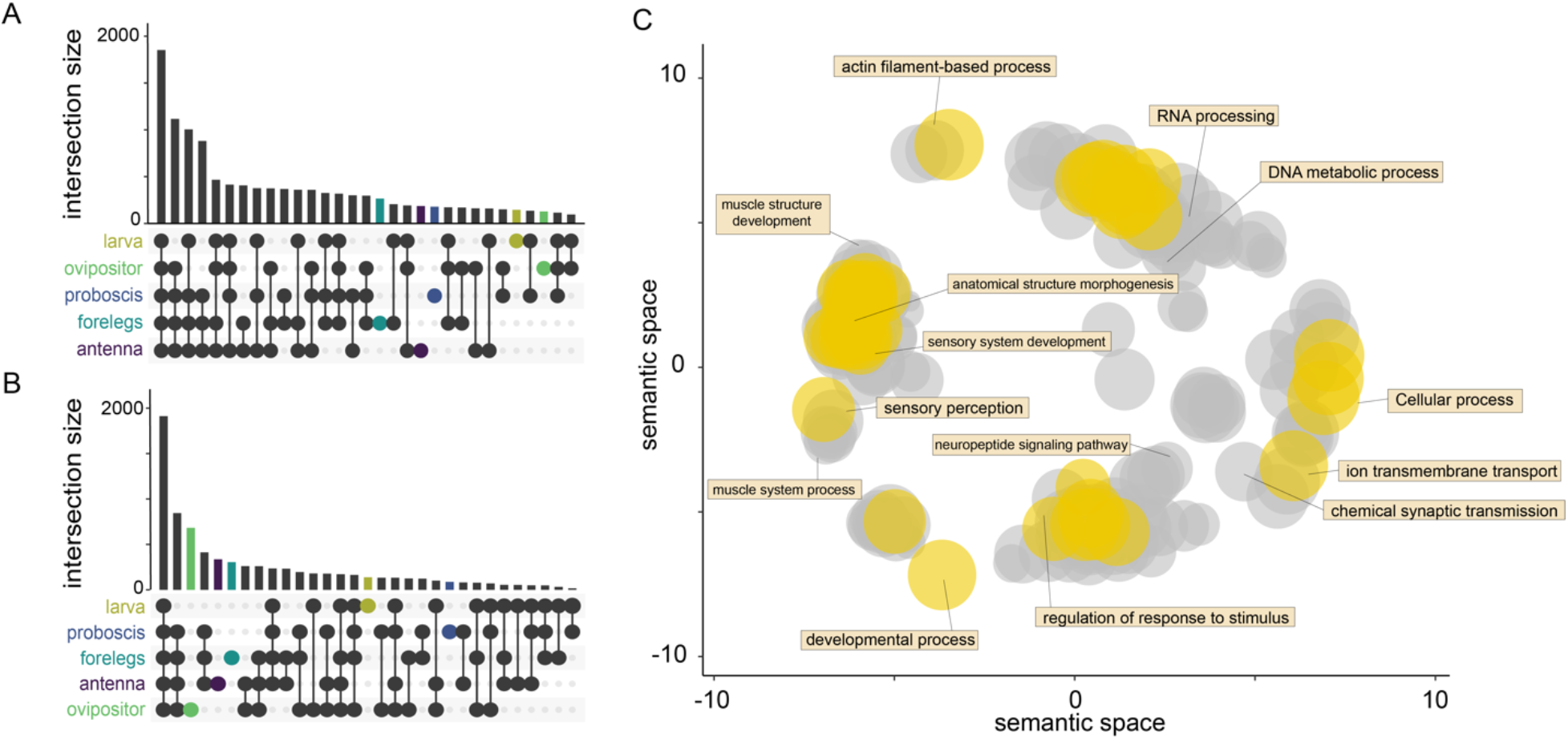
Comparison of the sets of differentially expressed genes identified by the phylogenetically-informed (OU) and DESeq2 methods. **(A)** The number of differentially expressed (DE) genes that overlap across tissue (regardless of species) using the l1ou method. Note, this is the same plot as in Fig. 2B and shows that most DE genes are shared across tissues. **(B)** The number of DE genes that overlap across tissues (regardless of species) using DESeq2 in pair-wise comparisons. DESeq2 gives comparable results to the l1ou method regarding the number of DE genes that are shared across tissues. **(C)** Multidimensional Scaling (MDS) from the two methods GO terms similarity matrix. Semantically similar GO terms project close to each other. Yellow circles show semantic terms shared between the two methods and gray circles show semantic terms unique to one or the other method. GO terms from the two methods clustered together indicating that DE genes from the two methods belong to similarly enriched pathways.

**Fig. S5.**
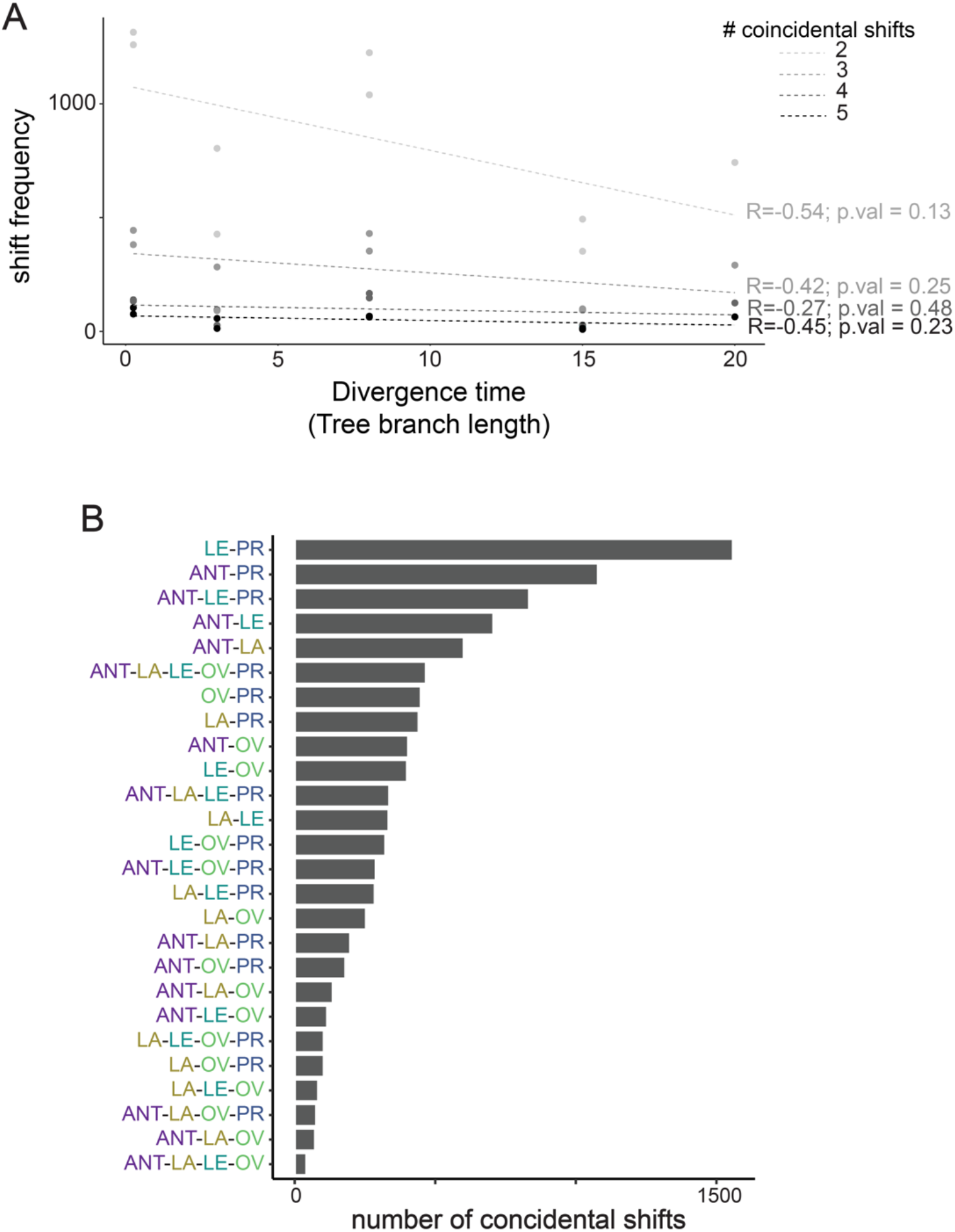
Investigation of coincidental shifts. **(A)** Number of coincidental shifts grouped by the number of coincidental changes and according to branch length (i.e. divergence time). The number of coincidental shifts are not correlated with branch length. **(B)** Frequency of coincidental shifts for all tissue combinations. Overall, the most frequent coincidental shift occurred between the legs and proboscis+palps.

**Fig. S6.**
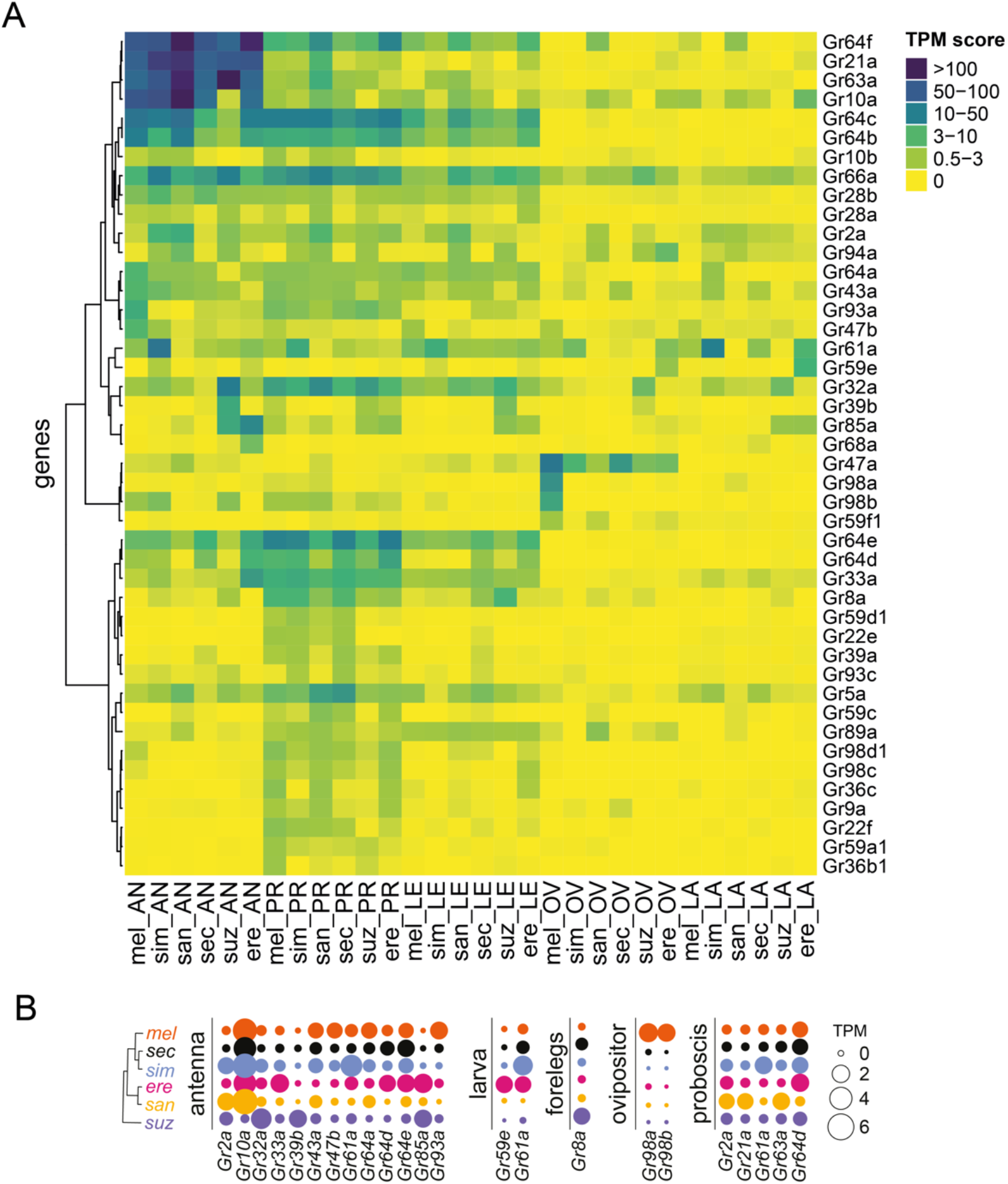
*Grs* expression across species and tissues. **(A)** Hierarchical clustering of mean *Gr* expression values (TPM). Each row contains a gene and each column contains a species’ tissue sample. Clustering was performed gene-wise. AN=antenna, PR=proboscis, LE=forelegs, OV= ovipositor, LA=larval head. **(B)** *Gr*s that have evolved species-specific expression gains or losses. Species names are abbreviated to the first three letters.

**Fig. S7.**
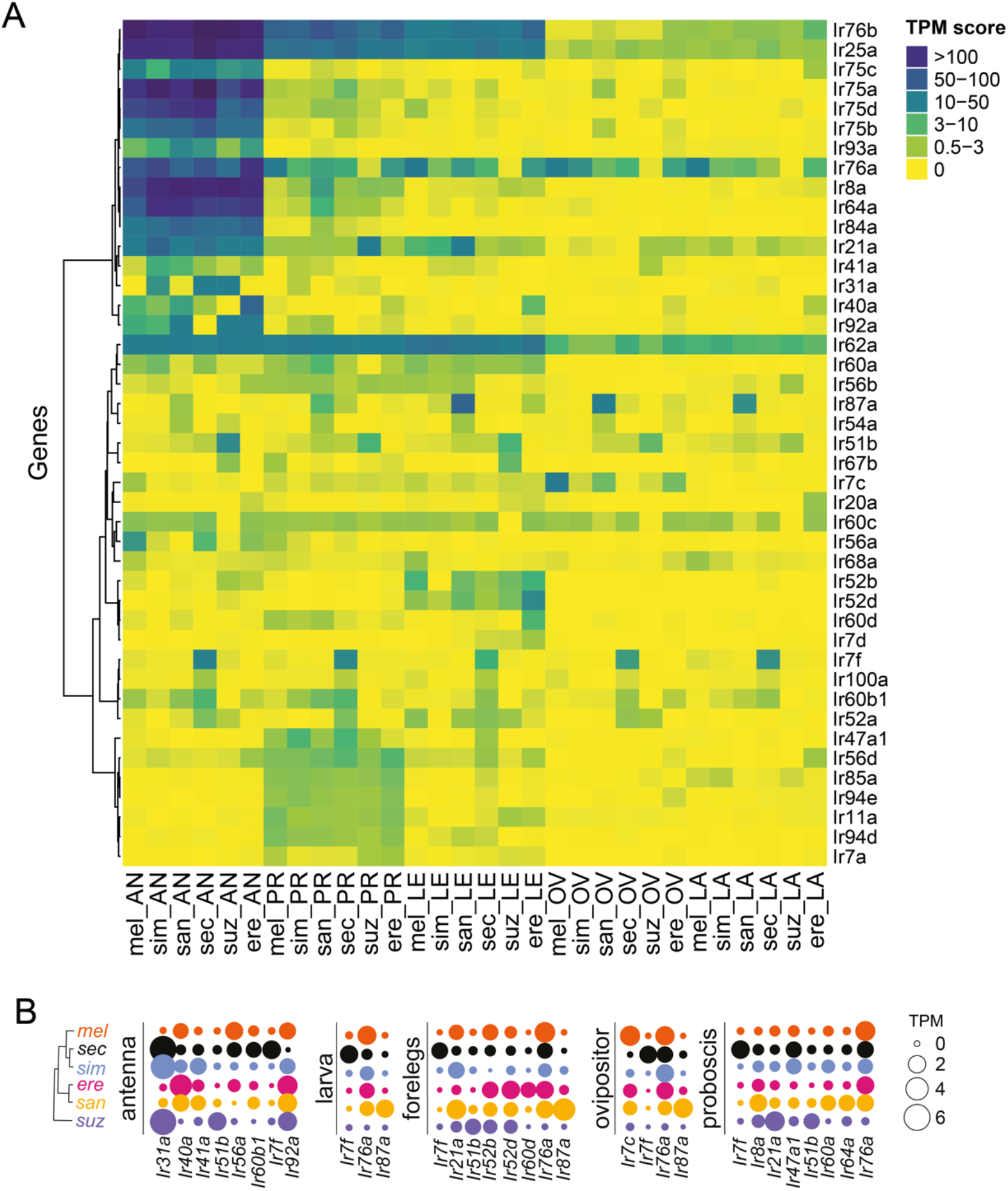
*Irs* expression across species and tissues. **(A)** Hierarchical clustering of mean *Ir* expression values (TPM). Each row represents a gene and each column a sample. Clustering was performed gene-wise. AN=antenna, PR=proboscis, LE=forelegs, OV= ovipositor, LA=larval head. **(B)** *Ir*s that have evolved species-specific expression gains or losses. Species names are abbreviated to the first three letters.

**Fig. S8.**
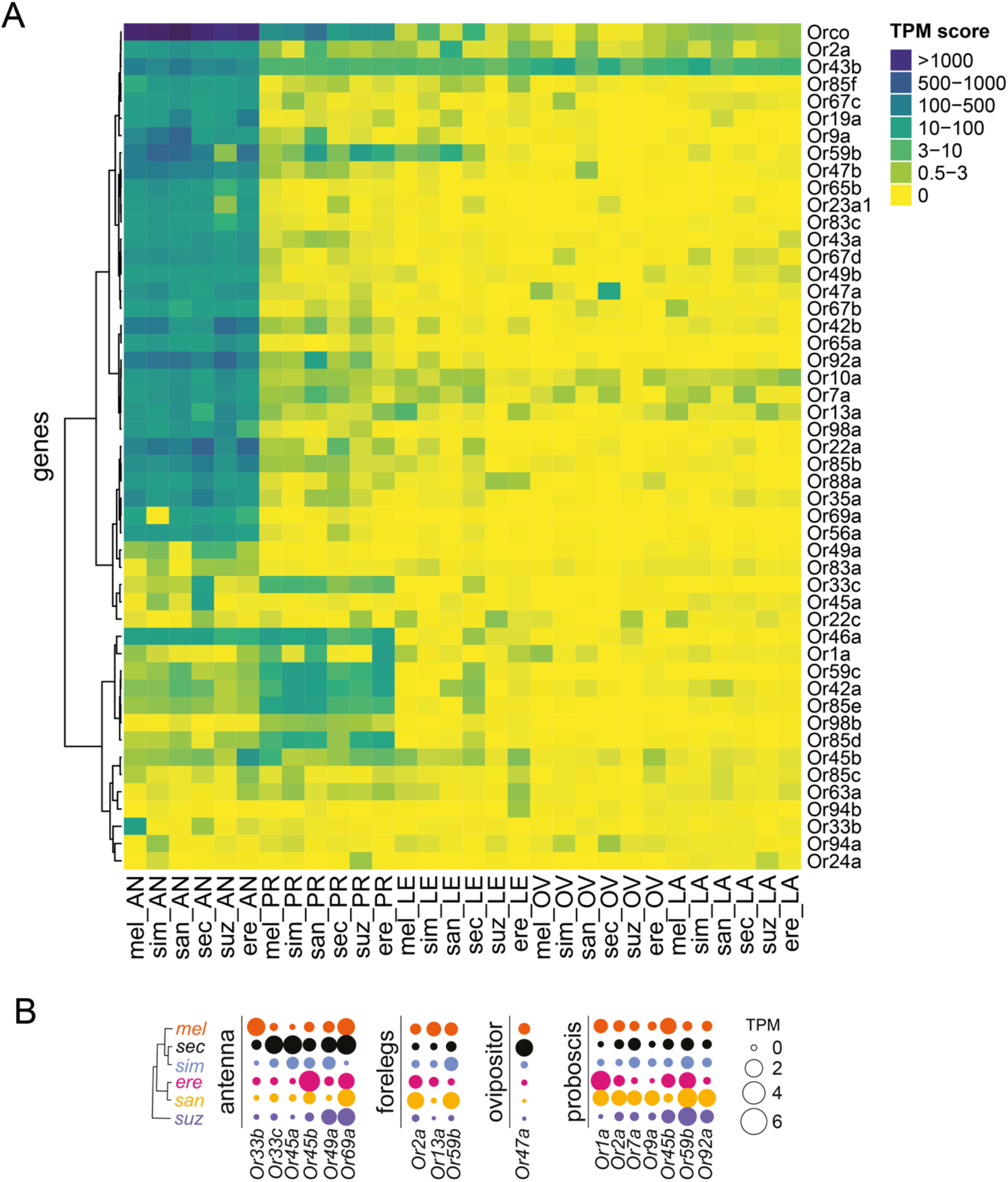
*Ors* expression across species and tissues. **(A)** Hierarchical clustering of mean *Or* expression values (TPM). Each row represents a gene and each column a sample. Clustering was performed gene-wise. AN=antenna, PR=proboscis, LE=forelegs, OV= ovipositor, LA=larval head. **(B)** *Ors* that have evolved species-specific expression gains or losses. Species names are abbreviated to the first three letters.

**Fig. S9.**
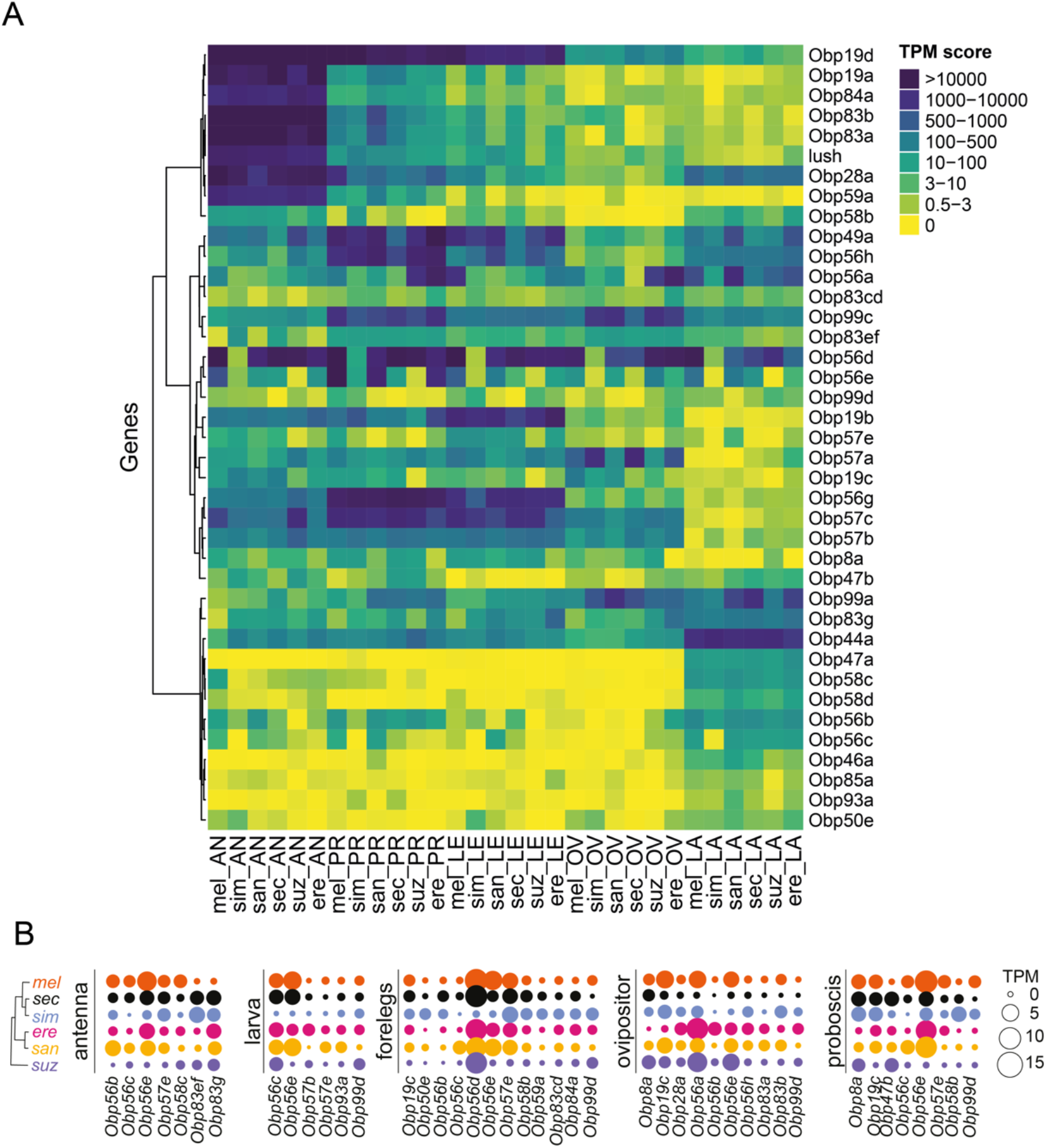
*Obp* expression across species and tissues. **(A)** Hierarchical clustering of mean *Obp* expression values (TPM). Each row represents a gene and each column a sample. Clustering was performed gene-wise. AN=antenna, PR=proboscis, LE=forelegs, OV= ovipositor, LA=larval head. **(B)** *Obp*s that have evolved species-specific expression gains or losses. Species names are abbreviated to the first three letters.

**Fig. S10.**
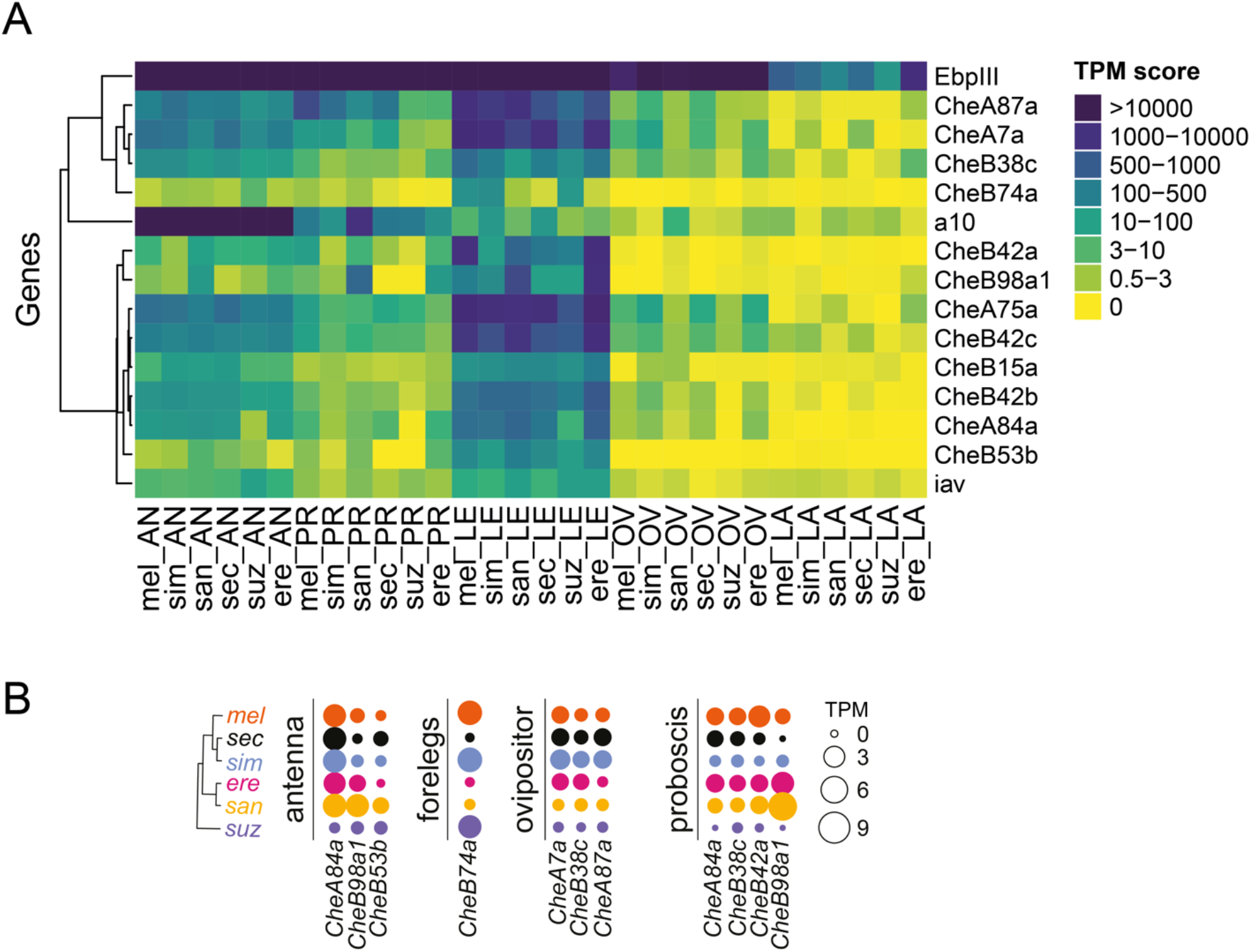
*CSPs* expression across species and tissues. **(A)** Hierarchical clustering of mean *CSP* expression values (TPM). Each row represents a gene and each column a sample. Clustering was performed gene-wise. AN=antenna, PR=proboscis, LE=forelegs, OV= ovipositor, LA=larval head. **(B)** *CSP*s that have evolved species-specific expression gains or losses. Species names are abbreviated to the first three letters

**Fig. S11.**
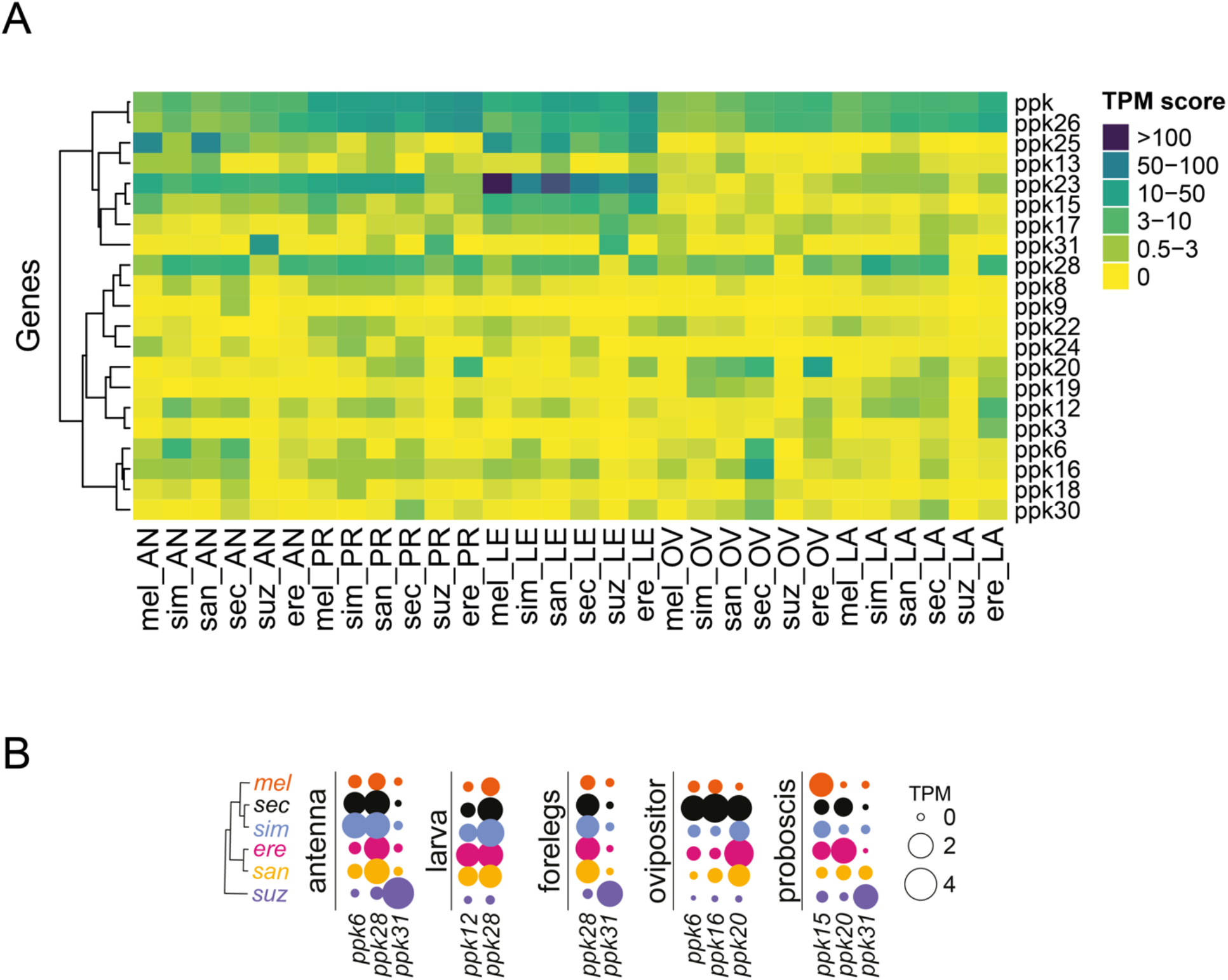
*Ppks* expression across species and tissues. **(A)** Hierarchical clustering of mean *ppk* expression values (TPM). Each row represents a gene and each column a sample. Clustering was performed gene-wise. AN=antenna, PR=proboscis, LE=forelegs, OV= ovipositor, LA=larval head. **(B)** *ppk*s that have evolved species-specific expression gains or losses. Species names are abbreviated to the first three letters

**Fig. S12.**
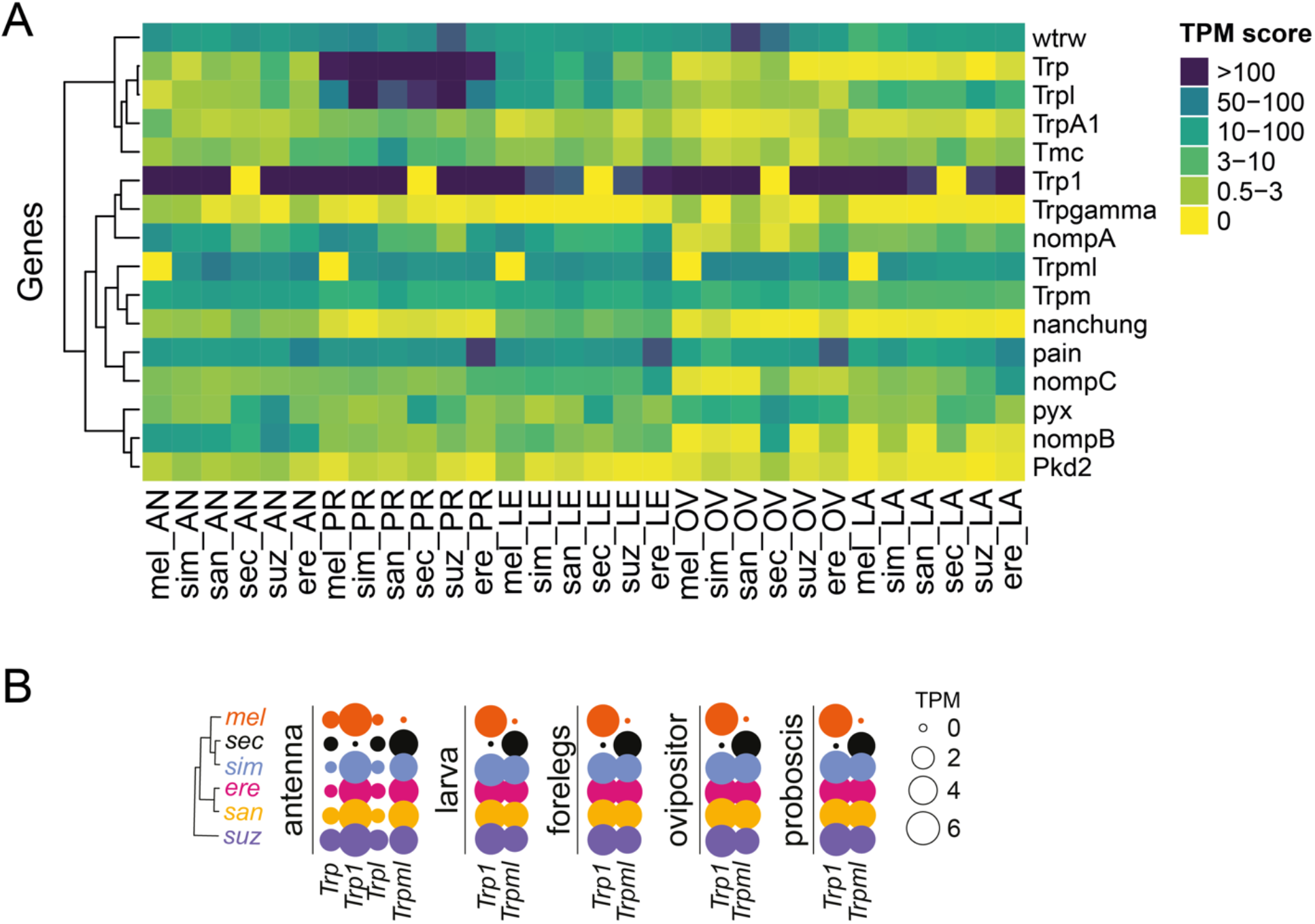
*Trp* expression across species and tissues. **(A)** Hierarchical clustering of mean *TRP* expression values (TPM). Each row represents a gene and each column a sample. Clustering was performed gene-wise. AN=antenna, PR=proboscis, LE=forelegs, OV= ovipositor, LA=larval head. **(B)** *TRP*s that have evolved species-specific expression gains or losses. Species names are abbreviated to the first three letters

**Fig. S13.**
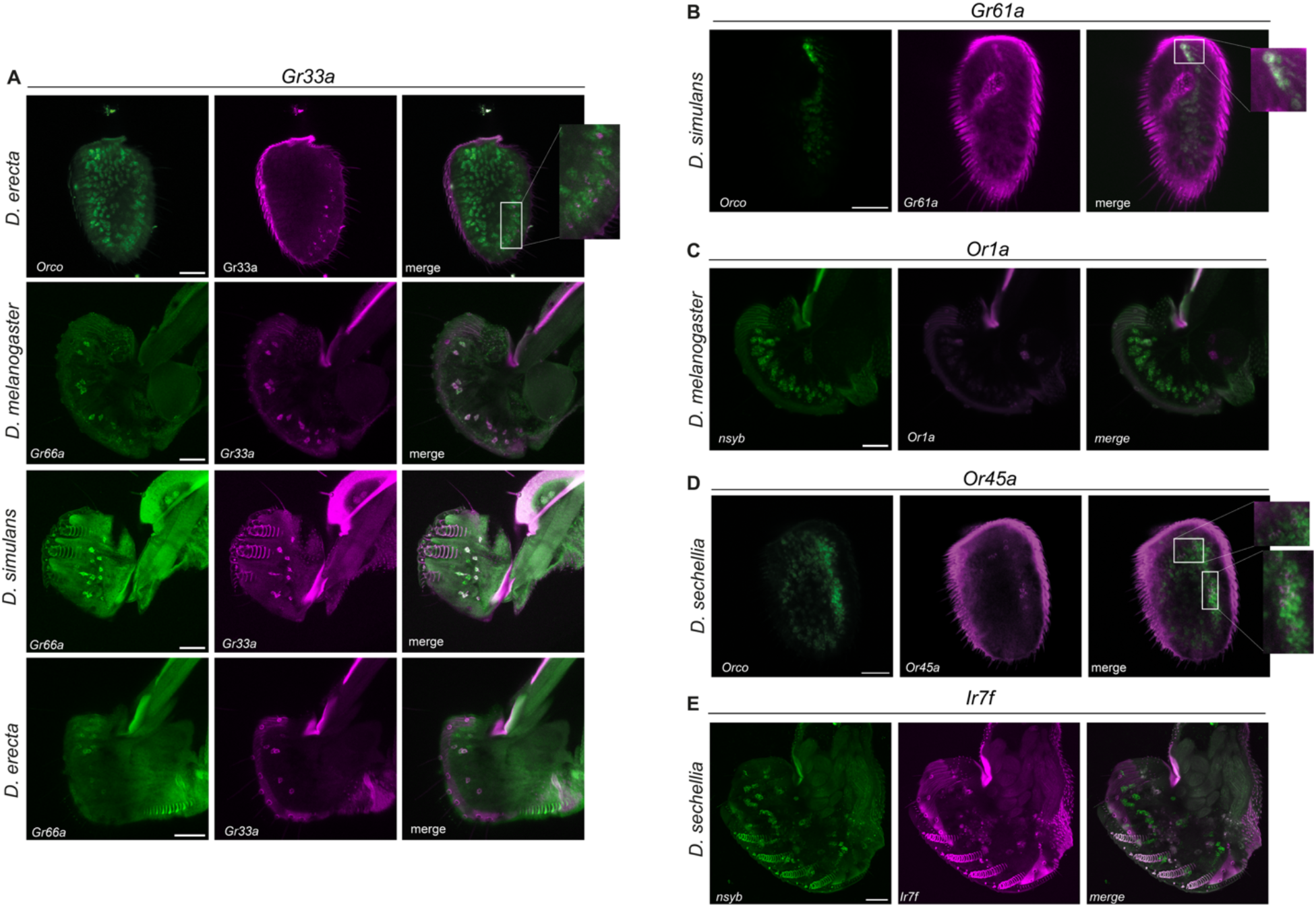
Expression analyses of chemosensory genes with cell markers. **(A)** Antennal specific gain of expression of *Gr33a* in *D.erecta*. Upper panel: RNA *in situ* hybridization of *Orco* and *Gr33a* in a whole mount antenna of *D. erecta*. Colabelled cells are magnified in the right panels. Bottom panels: RNA *in situ* hybridization of *Gr66a* and *Gr33a* in the labellum of *D. melanogaster*, *D. simulans* and *D. erecta*. We observed a conserved RNA-seq expression of *Gr33a* in the legs and proboscis samples across the six species but also found a high level of expression in *D. erecta*’s antenna. Consistent with our RNA-seq data, we were able to detect the expression of *Gr33a* in *D. erecta* antenna (17±4 cells) but not in the other two species. The *D. erecta*-specific antennal signal was verified by the successful detection of *Gr33a* in the proboscis of *D. simulans*, *D. sechellia*, and *D. melanogaster*, where *Gr33a* and *Gr66a* are co-expressed in bitter sensing neurons as previously described^1^. A co-labeling experiment using an *Orco* probe, which labels all olfactory sensory neurons, revealed *Gr33a*-*Orco* co-expression in the antenna, indicating that that *D. erecta*’s *Gr33a* has gained olfactory sensory neuron expression while maintaining its role in bitter taste sensing. **(B)** RNA in situ hybridization of *Orco* and *Or45a* in a whole mount antenna of *D. sechellia*. Co-labelled cells are magnified in the right panels. The expression of *Or45a* has been described as larva-specific in *D. melanogaster* and involved in aversive behavior to some volatile odors^2, 3^. While we did not detect *Or45a* in our *D. melanogaster* larval head RNA-seq datasets, likely due to low expression and/or the small number of cells that express it (mean TPM = 0.05), we did observe a high level of expression specifically in *D. sechellia* antenna (mean TPM = 20.25). By *in situ*, we were able to detect the expression *of Or45a in D. sechellia*’s antennal olfactory sensory neurons (11±4 cells) but not in *D. simulans* or *D. melanogaster* antenna. **(C)** RNA *in situ* hybridization of *nsyb* and *Ir7f* in the proboscis of *D. sechellia*. Ir7f belongs to a cluster of 7 tandemly arrayed paralogous Ionotropic receptors^8^. This cluster of Irs have diverse expression patterns in adults and larva, with *Ir7f* expression described as being specific to the dorsal pharyngeal sense organ in *D. melanogaster* larva^9^. Recently, a member of this clade, Ir7a, was shown to be an acetic acid sensor^10^, suggesting that the other Ir7 paralogs may recognize additional acids, though no further functional data exist for them. We were unable to detect an appreciable level of expression for *Ir7f* in the larva of the six species (TPM<0.3) but found lineage-specific gain of *Ir7f* expression for the other tissues in *D. sechellia*. We were able to co-labelled *Ir7f* and *nsyb* in *D.sechellia* labial palps, indicating its neuronal expression. Together our expression quantifications and cell detection suggest that *D. sechellia Ir7f* may play a specific function for this species which is an extreme host specialist that resides on noni fruit^11^. **(D)** RNA *in situ* hybridization of *Or1a* and *nsyb* in the labellum of *D. melanogaster*. *Or1a* has been described as larva-specific in *D. melanogaster* and involved in attraction to some odors^4^. Within our dataset, we found a high level of expression in the proboscis of *D. erecta* (mean TPM = 24.05) and appreciable levels of expression in the proboscis of *D. melanogaster and D. santomea* (mean TPM = 2.78 and 5.72 respectively). Consistent with these results, HCR-FISH probes for *Or1a* highlighted its expression in the proboscis of the three species but not in *D. simulans* (see Fig. 4D). Interestingly, the *Or1a* probe labeled cells larger than chemosensory neurons and located in a region that has not been described to contain sensory cells. To test if these cells belong to an unexpected population of neurons, we carried out a co-labelling experiment in *D. melanogaster* using the *Or1a* probe and a probe for a pan-neuronal marker, *neuronal Synaptobrevin* (*nSyb*). We detected broad *nSyb* expression throughout the proboscis but no co-localization with the *Or1a* probe. This result indicates that *Or1a* has evolved a non-neuronal expression in *D. erecta, D.santomea* and *D. melanogaster*. Based on their cuboidal morphology, we hypothesized that cells expressing *Or1a* are part of the salivary tract. **(E)** RNA *in situ* hybridization of *Orco* and *Gr61a* in *D. simulans* antenna. Co-labelled cells are magnified in the right panels. *Gr61a* has been identified as a glucose receptor in *D. melanogaster* and is expressed in neurons within the labellum, tarsal leg segments, and the labral sense organ^5–7^. Consistent with these descriptions, we have detected expression of *Gr61a* in these same tissues in *D. simulans* and *D. melanogaster* (mean TPM respectively in the labellum: 7.4 and 0.60, in the front legs: 5.3 and 2.13), in the legs of *D. santomea*, *D. sechellia*, *D.erecta* and *D.suzukii* (mean TPM= 0.67, 0.62, 1.39 and 0.82 respectively) and additionally in the antenna of *D. simulans*, *D. melanogaster*, *D. suzukii* and *D. erecta* (mean TPM= 23, 1.25, 0.53 and 1.49 respectively). As observed, the *Gr61a* expression was higher in all the *D. simulans*’ tissues than in the other species. A *D. simulans*-specific *in situ* probe confirmed the antennal olfactory sensory neurons expression of *Gr61a*. Though the signal for this probe was weak, we estimated ∼6 *Gr61a*-expressing cells.

1. Moon, S. J., Lee, Y., Jiao, Y. & Montell, C. A Drosophila gustatory receptor essential for aversive taste and inhibiting male-to-male courtship. Curr Biol 19, 1623–7 (2009).
2. Kreher, S. A., Mathew, D., Kim, J. & Carlson, J. R. Translation of Sensory Input into Behavioral Output via an Olfactory System. Neuron 59, 110–124 (2008).
3. Bellmann, D. et al. Optogenetically induced olfactory stimulation in Drosophila larvae reveales the neuronal basis of odor-aversion behavior. Front. Behav. Neurosci. 4, (2010).
4. Fishilevich, E. et al. Chemotaxis behavior mediated by single larval olfactory neurons in Drosophila. Curr Biol 15, 2086–96 (2005).
5. Kohatsu, S., Tanabe, N., Yamamoto, D. & Isono, K. Which Sugar to Take and How Much to Take? Two Distinct Decisions Mediated by Separate Sensory Channels. Front. Mol. Neurosci. 15, (2022).
6. Dahanukar, A., Lei, Y.-T., Kwon, J. Y. & Carlson, J. R. Two Gr Genes Underlie Sugar Reception in Drosophila. Neuron 56, 503–516 (2007).
7. Miyamoto, T., Chen, Y., Slone, J. & Amrein, H. Identification of a Drosophila Glucose Receptor Using Ca2+ Imaging of Single Chemosensory Neurons. PLOS ONE 8, e56304 (2013).
8. Croset, V. et al. Ancient Protostome Origin of Chemosensory Ionotropic Glutamate Receptors and the Evolution of Insect Taste and Olfaction. PLoS Genet. 6, e1001064 (2010).
9. Sánchez-Alcañiz, J. A. et al. An expression atlas of variant ionotropic glutamate receptors identifies a molecular basis of carbonation sensing. Nat. Commun. 9, 4252 (2018).
10. Rimal, S. et al. Mechanism of Acetic Acid Gustatory Repulsion in Drosophila. Cell Rep. 26, 1432-1442.e4 (2019).
11. R’Kha, S., Capy, P. & David, J. R. Host-plant specialization in the Drosophila melanogaster species complex: a physiological, behavioral, and genetical analysis. Proc. Natl. Acad. Sci. 88, 1835–1839 (1991). Neuron 46, 445–456 (2005).

**Fig. S14.**
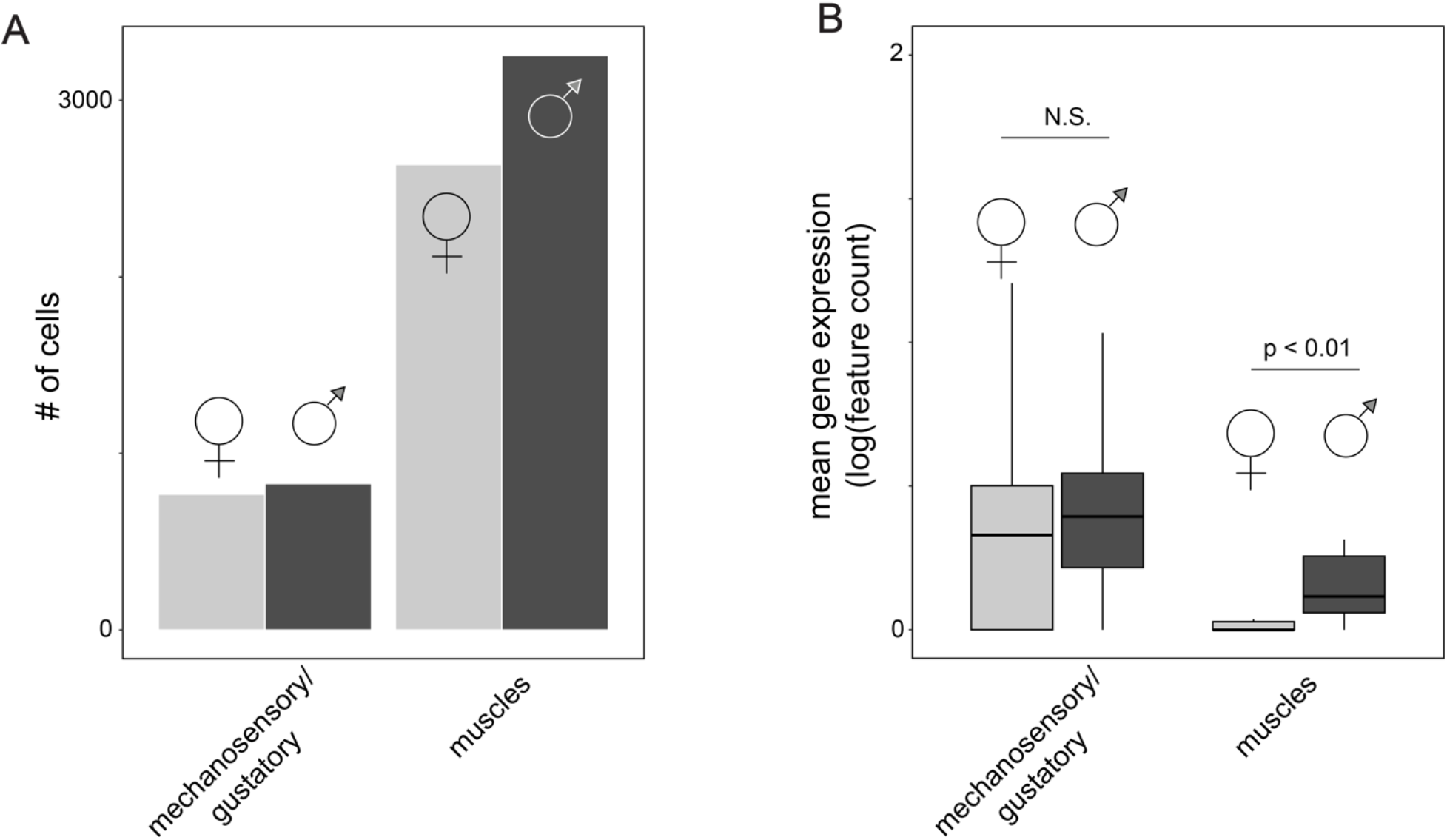
Comparison of male and female leg atlases. (A) Number of female and male cells in cell clusters enriched in male-biased genes from the Fly Cell Atlas dataset. (B) Boxplot of mean expression for male-biased genes in female (light gray) and male (dark gray) cells. Significance threshold from a Wilcoxon test between male and female mean expression are displayed.

## Supplemental Tables

**Table S1:** Differentially expressed genes based on all 1:1 orthologs. This table contains the branch and tissue that the expression change occurred in.

**Table S2:** List of chemosensory genes analyzed and the gene families that they belong to.

**Table S3:** Orthologous Chemosensory genes expression results compared with previous descriptions literature. Genes with a TPM threshold > 0,5 were defined as expressed. Each sheet is dedicated to a tissue. The “Summary” sheet presents the overall summary for all of the tissues’ comparisons. The “References” sheet provides the full list of papers used to carry out the comparisons.

*Colored rows indicate the following:*

green: 1:1 ortholog that has species-specific expression

blue: genes found expressed in our datasets but not in the literature,

yellow: agreement with tissues expression but different species (possible polymorphism) red: no match

purple: species-specific paralog

*Values within cells indicate the following:*

“ND”: denotes genes for which we could not find any evidence of expression in the literature. “N”: denotes a gene is absent in a given species.

**Table S4:** List of genes that were found to be expressed for an individual tissue across all species.

**Table S5:** Differentially expressed genes based on all the curated chemosensory gene sets of 1:1 orthologs. This table contains the branch the shift occurred and the tissues involved.

**Table S6:** Sex-biased genes separated by species and tissues.

**Table S7:** Detailed information on the molecular reagents used in this project.

**Table S8:** Lookup table connecting Flybase gene IDs to the gene symbols outputted by BRAKER.

## Supplemental Files

**File S1:** GTFs outputted by BRAKER.

**File S2:** RNA-seq Count tables for the full dataset generated by HTseq.

**File S3:** TPM count tables based on File S2 for all 1:1 orthologs.

**File S4:** GTF files based on conserved genic regions between the six species’ (“trimmed” GTFs) for the set of 1:1 orthologs.

**File S5:** RNA-seq Count tables based on the “trimmed” GTF file (File S4) generated by HTseq.

**File S6:** GTF files for the manually curated set of chemosensory genes.

**File S7:** TPM count tables for the manually annotated chemosensory gene set.

